# Contrasting synaptic roles of MDGA1 and MDGA2

**DOI:** 10.1101/2023.05.25.542333

**Authors:** Michael A. Bemben, Matthew Sandoval, Aliza A. Le, Sehoon Won, Vivian N. Chau, Julie C. Lauterborn, Salvatore Incontro, Kathy H. Li, Alma L. Burlingame, Katherine W. Roche, Christine M. Gall, Roger A. Nicoll, Javier Diaz-Alonso

**Affiliations:** Department of Cellular and Molecular Pharmacology, University of California at San Francisco, San Francisco, CA 94158, USA; Department of Anatomy & Neurobiology, University of California at Irvine, CA, 92617, USA; Center for the Neurobiology of Learning and Memory, University of California at Irvine, CA, USA; Receptor Biology Section, National Institute of Neurological Disorders and Stroke (NINDS), National Institutes of Health (NIH), Bethesda, MD, 20892, USA; Unité de Neurobiologie des canaux Ioniques et de la Synapse (UNIS), UMR1072, INSERM, Aix-Marseille University, Marseille, 13015, France; Department of Pharmaceutical Chemistry, University of California, San Francisco, San Francisco, CA 94158, USA; Department of Physiology, University of California at San Francisco, San Francisco, CA, 94158, USA

## Abstract

Neurodevelopmental disorders are frequently linked to mutations in synaptic organizing molecules. MAM domain containing glycosylphosphatidylinositol anchor 1 and 2 (MDGA1 and MDGA2) are a family of synaptic organizers suggested to play an unusual role as synaptic repressors, but studies offer conflicting evidence for their localization. Using epitope-tagged MDGA1 and MDGA2 knock-in mice, we found that native MDGAs are expressed throughout the brain, peaking early in postnatal development. Surprisingly, endogenous MDGA1 was enriched at excitatory, but not inhibitory, synapses. Both shRNA knockdown and CRISPR/Cas9 knockout of MDGA1 resulted in cell-autonomous, specific impairment of AMPA receptor-mediated synaptic transmission, without affecting GABAergic transmission. Conversely, MDGA2 knockdown/knockout selectively depressed NMDA receptor-mediated transmission but *enhanced* inhibitory transmission. Our results establish that MDGA2 acts as a synaptic repressor, but only at inhibitory synapses, whereas both MDGAs are required for excitatory transmission. This nonoverlapping division of labor between two highly conserved synaptic proteins is unprecedented.

**Teaser:** MDGAs 1 and 2 independently localize to and modulate excitatory and inhibitory hippocampal synapses by different mechanisms.

## Introduction

The brain integrates and processes information via cell-to-cell communication at specializations between neurons termed synapses. Synapses link an individual neuron to a complex network of interconnected cells and facilitate the transfer of information in the form of excitation and inhibition. The balance and integration of excitatory and inhibitory synaptic transmission is required for proper brain function, as disruptions in these processes lead to neurological disorders such as epilepsy, autism spectrum disorders (ASDs), and schizophrenia (*1–3*). The assembly, maturation and maintenance of synapses are sustained by a multifarious network of proteins that organize and align the presynaptic release and postsynaptic receptor sites to allow for effective communication between neurons (*4–6*).

Similar to genetic alterations in other synaptic proteins that affect function, mutations in the memprin, A5 protein, receptor protein tyrosine phosphatase mu (MAM) domain containing glycosylphosphatidylinositol anchor (MDGA) family of proteins have been implicated in cognitive and psychiatric disorders, underscoring their critical importance in brain function (*7–9*). MDGAs are membrane-associated proteins that contain six tandem immunoglobulin (Ig)-like domains, a fibronectin-like region (FNIII), a single MAM domain, and a C-terminal glycosylphosphatidylinositol (GPI) anchor [Fig. 1A, B (*10*)]. The expression of MDGA proteins is restricted to the nervous system, begins early in development and continues throughout adulthood (*11*). Mammals have two highly conserved MDGAs, MDGA1 and MDGA2, which their dysfunction are associated, with schizophrenia and ASDs, respectively. (*7–9*). There is debate regarding the type of synapses each isoform localizes to due to lack of reliable anti-MDGA antibodies, inconclusive results in recombinantly expressed MDGA studies and inconsistency in the interpretation of knockdown experiments. Overexpressed YFP-MDGA1 and HRP-MDGA1 localizes to both excitatory and inhibitory synapses, as well as to extrasynaptic sites (*12, 13*). Similarly, HRP-tagged MDGA2 was found at inhibitory synapses (*13*) with no obvious concentration at excitatory, inhibitory, or extrasynaptic sites in another study (*14*).

**Fig 1.**
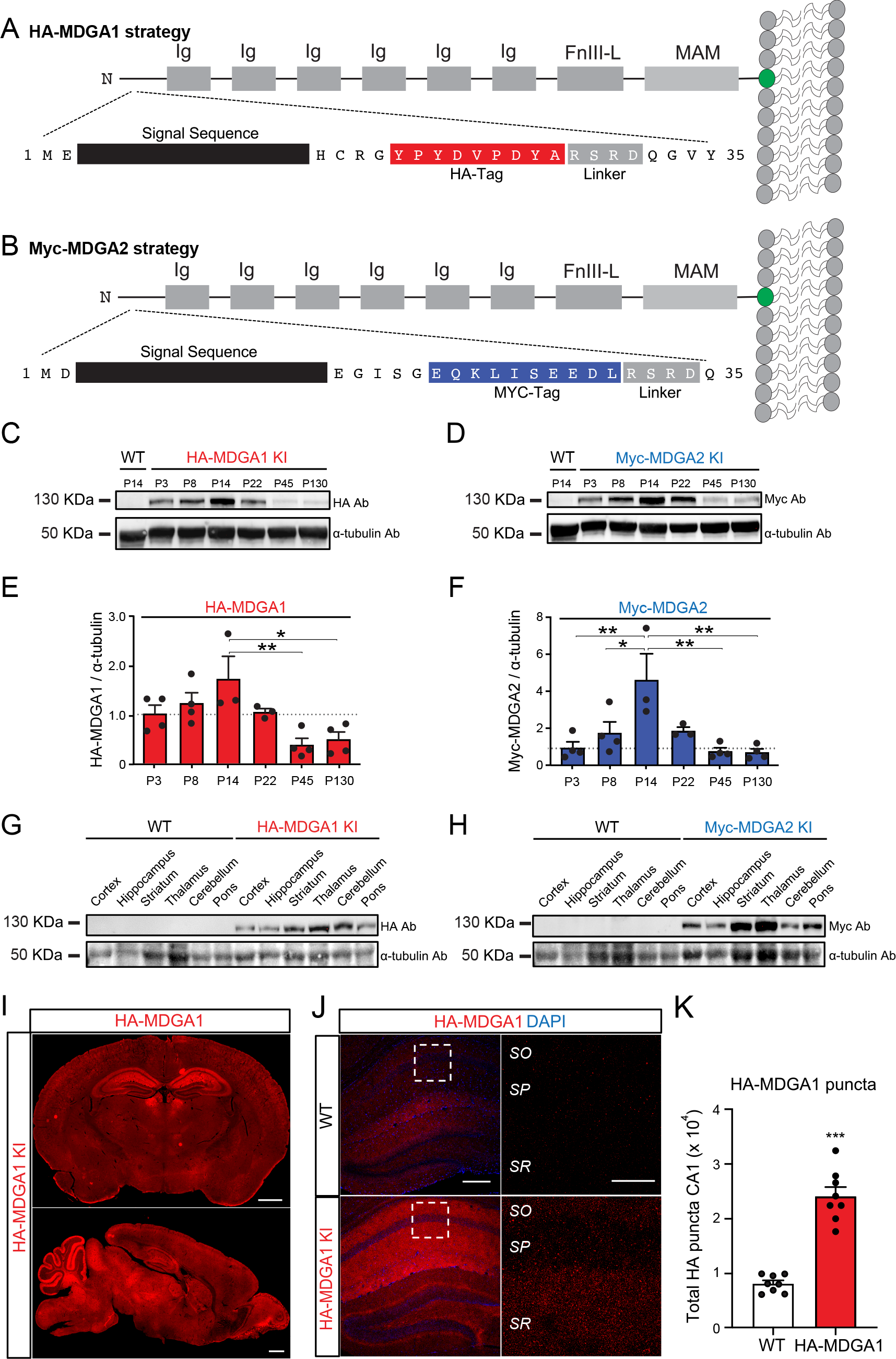
Native MDGA1 and MDGA2 expression in the postnatal mouse brain. A, B, Schematic of the epitope-tagged HA-MDGA1 (A) and Myc-MDGA2 (B). C, D, Representative immunoblot of HA-MDGA1 and Myc-MDGA2 expression in the mouse brain, respectively, from postnatal day 3 to postnatal day 130, using α-tubulin as a loading control. WT P14 sample included for antibody specificity control. E, F, Quantification of HA-MDGA1 / α-tubulin and Myc-MDGA2 / α-tubulin ratios, respectively, normalized to postnatal day 3 values in the KI mice. P14 (peak HA-MDGA1 and Myc-MDGA2 expression) samples from WT mice were used as a control of tag antibody specificity. G, H, Representative immunoblots of HA-MDGA1 and Myc-MDGA2, respectively, using α-tubulin as a loading control, across brain regions. n=3-4 mice/condition for all Western blot analyses. I, Representative HA- MDGA1 immunofluorescence staining in coronal (top panel) and sagittal (bottom panel) slices of HA-MDGA1 mice at P15. J, Representative confocal images of HA staining with DAPI in WT (top) and HA-MDGA1 (bottom) of P20 HA-MDGA1 KI mouse hippocampus. High magnification insets of area CA1 are shown to the right. K, Quantification of HA puncta from high magnification confocal images shown in J (p<0.001, n=8 mice/genotype). Bar graphs represent mean ± SEM. *, p<0.005; **, p<0.01; ***, p<0.001, one-way ANOVA (E, F), unpaired two-tailed Student’s T-Test (K). Scale bars: I, 1 mm. J, 200 µm, insets: 50 µm. “SR”, stratum radiatum; “SP”, stratum pyramidale; “SO”, stratum oriens.

Similarly, a recent study found that expressed epitope-tagged MDGAs are highly mobile in the plasma membrane, with only a small fraction localizing at synapses (*15*). Despite these localization studies, functional readouts from overexpression studies have presumed MDGA localization to postsynaptic sites (see discussion below). In stark contrast with overexpression studies, an elegant, *in situ* proximity-based proteomic characterization found endogenous MDGA1 and MDGA2 localized to excitatory and inhibitory postsynapses, respectively (*13*). Success in generating truly specific antibodies against MDGAs has been limited (*16*). However, a recent study reported the generation of a specific antibody against MDGA1 (*15*) and found a higher proportion of MDGA1 at excitatory synapses (40-45%) *vs* inhibitory synapses (20%) *in vitro*. Puzzlingly, this preference disappeared with neuronal culture maturation. Overall, these findings suggest that overexpressed and endogenous MDGAs localize differently, which is not uncommon for synaptic cell adhesion molecules.

Despite the lack of definitive subcellular localization, exogenous MDGA1 expression in cultured neurons has been shown to decrease inhibitory synapse number (*12, 17, 18*), whereas MDGA2 expression decreases excitatory and inhibitory synapse number and transmission (*14*). These results imply that MDGA1 is localized to inhibitory synapses, whereas MDGA2 is localized to both excitatory and inhibitory synapses, in contrast with reported localization of endogenous MDGAs (Loh *et al.*, 2016). But, most importantly, these results indicate that MDGAs are synaptic repressors, an unusual role for synaptic proteins. Consistent with this model, i) shRNA-mediated MDGA1knockdown (KD) increases inhibitory synapse density and transmission in cultured neurons (*12, 13, 17*) – although this has been recently challenged by studies which find no specific change in inhibitory synaptic transmission upon MDGA1 deletion in cultured and hippocampal CA1 neurons (*15, 18*), and ii) MDGA2 KD/knockout (KO) leads to a specific increase in excitatory transmission (*15*) – although another group found that it does not result in a specific change in either excitatory or inhibitory synapse number (*13*). MDGA1 KO mice are viable with no gross anatomical phenotypes, but have an imbalance of excitation/inhibition (*19, 20*). In contrast, the MDGA2 KO is lethal in mice (a striking phenotype for an individual synaptic protein), which has hampered its study (*14*).

MDGA’s synaptic repressor function is thought to rely on their ability to modulate transsynaptic neuroligin-neurexin interactions, thereby interfering with a core molecular substrate of synapse formation and maintenance (*4, 16, 21*). Consistent with cell adhesion and surface bindings assays (*12, 17*), multiple independent groups have provided robust MDGA –neuroligin co-crystal structural data suggesting that MDGAs can sterically block access of neurexins to neuroligins (*22–24*). However, this model is primarily based on exogenous overexpression of MDGAs, and despite the reported high affinity of the interaction between MDGA1 and neuroligin-2 (Nlgn2), it remains unclear whether these proteins share overlapping spatial or temporal expression to interact *in vivo* (*25*). It is noteworthy that individual neuroligin KOs are not lethal, suggesting that MDGA2, at least in part, performs neuroligin-independent functions. Finally, unbiased proteomic screens have not identified MDGAs as binders to neuroligin, also challenging the model (*26*).

In summary, previous findings suggest that MDGAs can act as synaptic repressors, and have at least partially non-overlapping expression in the brain, but there is not a single unified model that can incorporate all of the field’s findings. Critically, the endogenous localization and role of MDGA proteins remains undefined. Here, we systematically address two fundamental and unresolved questions in the field: i) are endogenous MDGAs primarily synaptic repressors acting through neuroligins? and ii) what are the endogenous (spatial and temporal) localization of MDGA1 and MDGA2? Our results show that MDGA1 and MDGA2 expression overlap temporally and spatially in the developing mouse brain. In agreement with its expected role as synaptic repressor, we found that loss of function of MDGA2 results in increased synaptic transmission, although, contrary to expectations, only at inhibitory synapses. Surprisingly, we found that both MDGA1 and MDGA2 are required for excitatory synaptic transmission, with an unprecedented segregation of tasks for a family of synaptic proteins: while MDGA1 contributes exclusively to AMPAR- mediated transmission, MDGA2 selectively supports NMDAR-mediated transmission. Using a combination of techniques, primarily focusing on endogenous MDGAs in the Schaffer collateral (SC)-CA1 pyramidal neuron (PN) synapse, we describe unrecognized roles for MDGA1 and MDGA2 controlling excitatory and inhibitory synaptic transmission by distinct mechanisms in the mouse hippocampus.

## Results

### Temporal and anatomical distribution of native MDGA proteins

Where are endogenous MDGA1 and MDGA2 expressed? To reliably assess the temporal and spatial expression patterns of endogenous MDGA1 and MDGA2, we generated epitope-tagged knock-in (KI) mouse lines. Specifically, we generated hemagglutinin (HA)-tagged MDGA1 mice (Fig. 1A, Fig. S1) and Myc-tagged MDGA2 mice (Fig. 1B, Fig. S2). A series of steps were taken to ensure consistency with the literature and to provide confidence that the tags would not alter the function of MDGAs: i) we inserted small tags (8-10AAs) in the N-terminal of the proteins to minimize the likelihood of altering protein folding, trafficking and/or function given that previous work with fluorescently tagged MDGAs showed widespread localization in dendrites and axons (*12*); ii) we specifically used HA-tag for MDGA1 and Myc-tag for MDGA2 in the same locations as the commonly used overexpression constructs in the field; and iii) throughout the manuscript we focused the characterization of native MDGAs at hippocampal CA1 PN, given that both MDGA1 and MDGA2 are expressed in CA1 PNs [Mouse Whole Cortex and Hippocampus SMART-seq [2019]) with 10x-SMART-seq taxonomy (2021), (*14, 19, 20*)]. First, we validated the specificity of the HA-MDGA1 and Myc-MDGA2 signal using Western blots [WB, (Fig. 1C-H)]. Bands were detected at approximately 130 KDa, slightly above the expected molecular weight of MDGA1 and MDGA2, yet consistent with recent reports and likely due to prominent glycosylation (*15*). These bands were absent from wild-type (WT) samples and considered specific. Using HA-MDGA1 / Myc-MDGA2 and WT mice, we characterized the expression of the MDGAs across postnatal developmental stages focusing on six different time points, from postnatal day 3 (P3) to P130. The expression of both native MDGAs is strongly developmentally regulated, with a peak around P14 and more modest expression extending into adulthood (Fig. 1C-F). We next assessed the regional distribution across different brain regions at P15. We detected high expression of MDGA1 in areas largely consistent with previous *in situ* hybridization data (*17*) and β-galactosidase activity in Mdga1^+/lacZ^ mice (*19*) (Fig. 1G). Myc- MDGA2 expression followed a similar pattern (Fig. 1H), also consistent with previous estimations of MDGA2 expression using β-galactosidase activity in Mdga2^+/lacZ^ mice (*14*).

Immunofluorescence confirmed that native MDGA1 is highly expressed in the areas identified with WB, displaying particularly high expression in the hippocampus (Fig. 1I-K).

### Synaptic localization of MDGA1

MDGA proteins are considered synaptic adhesion molecules (*16, 25, 27*). However, while some reports have provided evidence of synaptic expression of endogenous MDGAs (*13*), single molecule imaging studies on overexpressed MDGAs suggest a diffuse localization within dendrites (*15*). Therefore, we set out to determine if MDGA1 is expressed at synapses, and whether it preferentially localizes to excitatory or inhibitory synapses. Based on the developmental time-course of MDGA1 expression (Fig. 1C, E), we initially co-labelled P15 brain samples for HA-MDGA1 together with excitatory or inhibitory synaptic markers. We focused on hippocampal area CA1, which shows high expression both at the mRNA (*17*) and protein levels (Fig. 1). Using confocal microscopy, we detected a punctate distribution of HA- MDGA1 in strata radiatum (SR) and oriens (SO), whereas the density of HA-MDGA1 puncta was minimal in stratum pyramidale [SP, (Fig. 1J, K)].

These findings indicate that native MDGA1 displays a punctate, synapse-like distribution in CA1, with a laminar distribution consistent with a bias towards excitatory synapses. To directly test this prediction, we evaluated MDGA1 localization in CA1 SR, which includes excitatory CA3 (SC)-CA1 PN synapses as well as inhibitory synapses impinging on dendritic shafts. To determine whether native MDGA1 localizes at excitator y synapses, we co-labeled P15 HA-MDGA1 and WT samples with HA and the excitatory postsynaptic marker Homer1b/c. Using fluorescence tomography based on three-dimensional (3D) reconstructions of individual HA-MDGA1 puncta alongside the excitatory post-synaptic puncta created from image z-stacks, we examined the colocalization of MDGA1 with Homer. We then performed a similar analysis with the postsynaptic inhibitory synaptic marker Nlgn2, previously shown to interact with MDGA1 (*12, 17, 22-24*), (Fig. S3A). Counts of the double labeled synapses indicate that native MDGA1 is enriched at excitatory Homer1b/c- immunoreactive (ir) postsynapses, compared with inhibitory Nlgn2-ir postsynapses (Fig. S3A, B). We then assessed colocalization of MDGA1 with markers for the presynaptic element at excitatory and inhibitory synapses, vGluT1 and vGAT, respectively. Quantification of double- labeled puncta again indicated a preferential localization of HA-MDGA1 to excitatory presynaptic elements (Fig. S3G, H). Despite the presence of some HA immunolabeling in WT samples, the number of HA-ir puncta was substantially higher in KI samples (Fig. S3C, E, I, K), whereas numbers of elements immunoreactive for synaptic markers did not differ between genotypes (Fig. S3D, F, J, L). We then performed 3D reconstructions using confocal z-stacks from the SR using P20 mice, which exhibit substantial MDGA1 expression and have more mature synapses, and quantified the number of HA-MDGA1 puncta overlapping with excitatory and inhibitory postsynaptic markers (Fig. 2A, B) and presynaptic markers (Fig. 2G, H). Using this approach, the number of HA-ir puncta was several-fold higher in HA-MDGA1 samples compared with WT counterparts (Fig. 2C, E, I, K), while the number of synaptic marker puncta was not significantly different between genotypes (Fig. 2D, F, J, L). Among the HA-MDGA1 puncta localized at postsynapses, we found a greater number of puncta localized in Homer1b/c-ir excitatory compartments, compared with Nlgn2-labelled inhibitory compartments (Fig. 2B). In contrast, we found a non-significantly higher number of MDGA1 associated with the excitatory presynaptic marker vGluT1 than with the inhibitory marker vGAT. Together, our findings indicate that native MDGA1 is preferentially expressed at excitatory synapses, although, despite the punctate pattern of expression found for native MDGA1, a substantial proportion of MDGA1 was not found to colocalize with the synaptic markers analyzed here.

**Fig 2.**
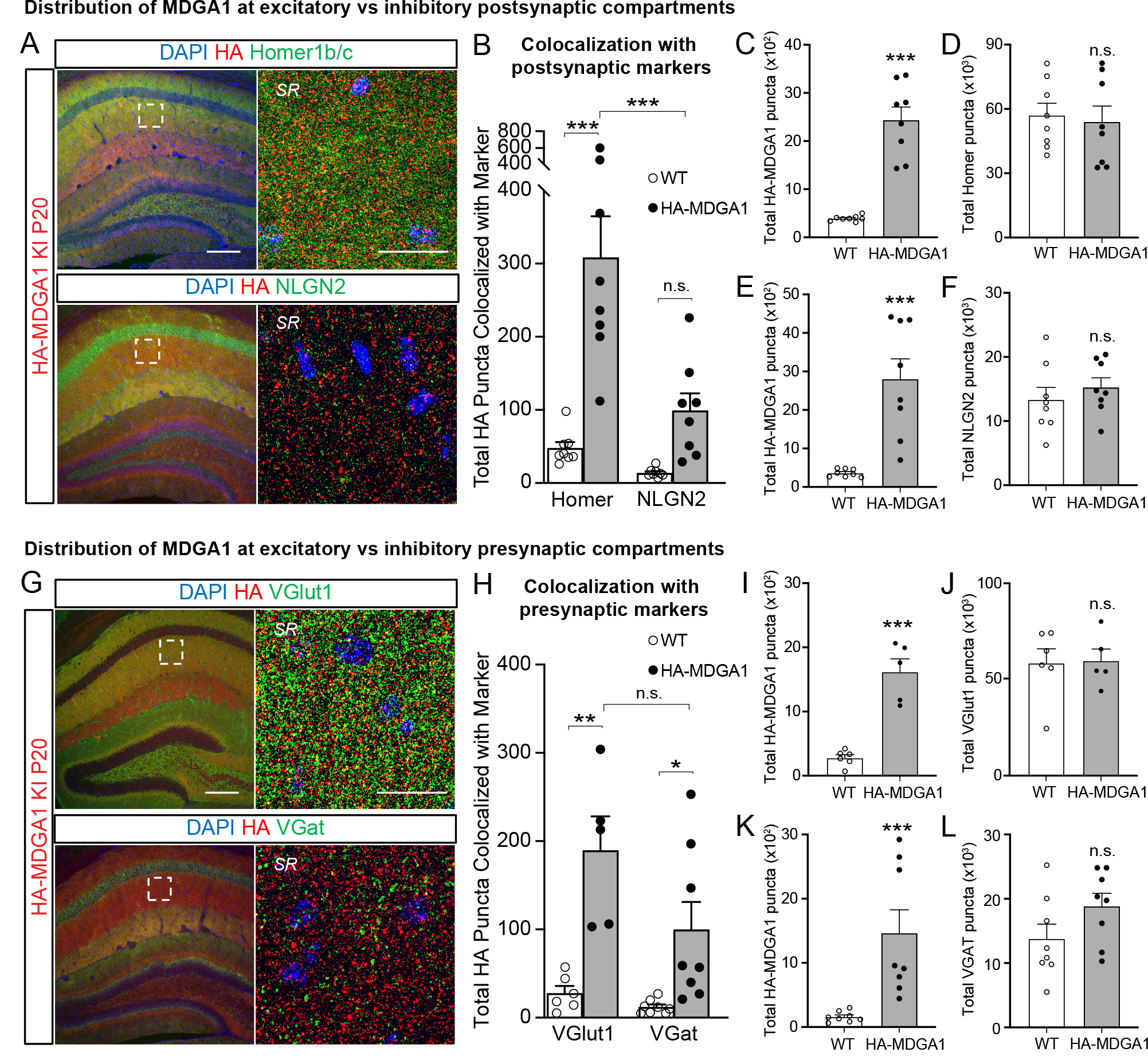
HA-MDGA1 is enriched in excitatory synaptic compartments in the stratum radiatum of the mouse hippocampal area CA1. A, Low magnification (left) and high magnification (right) representative confocal immunofluorescence images labeling DAPI, HA-MDGA1, and either excitatory (Homer1b/c, top panels) or inhibitory (Nlgn2, bottom panels) postsynaptic marker in P20 HA-MDGA1 mouse hippocampi. B, Quantification of the colocalization of high intensity HA- MDGA1 and synaptic marker puncta (within 5 μm of each other) in the SR. The number of colocalized HA puncta is higher in HA-MDGA1 than WT with Homer1b/c (p <0.0001), but not with Nlgn2 p = 0.2219). In HA-MDGA1 samples, the proportion of HA-MDGA1 puncta colocalized with Homer1b/c is higher than with Nlgn2 (p< 0.0003), whereas in WT samples it is not (p=0.9981). C, D, In the MDGA1-HA/Homer1b/c colocalization experiment, the number of HA puncta was higher in KI samples (p<0.0001, C), but the number of Homer1b/c was not different (p*=*0.7481, D). E, F, in the HA-MDGA1/Nlgn2 colocalization experiment, the number of HA puncta was higher in KI samples (p=0.0004, E), but the number of Nlgn2 was not (p*=*0.4332, F). G, Low magnification (left) and high magnification (right) representative confocal immunofluorescence images labeling DAPI, HA-MDGA1, and either excitatory (vGluT1, top panels) or inhibitory (vGAT, bottom panels) presynaptic marker in P20 HA-MDGA1 mouse hippocampi. H, Quantification of the colocalization of high intensity HA-MDGA1 and synaptic marker puncta (within 5 μm of each other) in the SR. The number of colocalized HA puncta is higher in HA-MDGA1 than WT (p = 0.0012 with vGluT1, p = 0.049 with vGAT). In neither HA- MDGA1 samples (p = 0.0958) nor in WT samples (p=0.9981) the proportion of HA-MDGA1 puncta colocalized with vGluT1 and vGAT are significantly different. I, J, In the MDGA1- HA/vGluT1 colocalization experiment, the number of HA puncta was higher in KI samples (p<0.0001, I), and the number of vGluT1 was not (p*=* 0.9033, J). K, L, in the HA- MDGA1/vGAT colocalization experiment, the number of HA puncta was higher in KI samples (p= 0.0029, K), but the number of vGAT was not (p= 0.1157, L). n = 5/8 mice/group. Bar graphs represent mean ± SEM. n.s., non-statistically significant; *, p < 0.05; **, p<0.01; ***, p<0.001, Two-way ANOVA followed Tukey’s post hoc test in B and H and unpaired T-test in C-F and I-L. Scale bars: Left panels: 200 µm. Right panels: 20 µm. “SR”, Stratum Radiatum.

### Synaptic localization of MDGA2

Our KI strategy enables MDGA2 protein detection via WB, allowing for detection of MDGA2 protein in mouse brain for the first time, which constitutes a significant advance in the field (Fig. 1D, F, H). Unfortunately, we were unable to identify conditions for Myc-MDGA2 localization using immunofluorescence. Thus, we turned to an unbiased proteomics screen to decipher the landscape surrounding MDGA2 to provide insights into its localization.

Immunoprecipitated MDGAs were exposed to solubilized mouse forebrain lysates to perform an unbiased proteomic analysis (Fig. 3A). Synaptic proteins were enriched with both MDGA1 and MDGA2. Consistent with our previous findings, excitatory synaptic proteins, such as Leucine-rich repeat transmembrane neuronal 1 (LRRTM1), are enriched in the MDGA1 proteome, whereas we identified proteins found in both excitatory [e.g., NMDA receptor (NMDAR) subunit GluN1], and inhibitory synapses (such as Nlgn2) in the MDGA2 proteome (Fig. 3B, supplementary Table 1 for a full list of MDGA1 and MDGA2 interacting proteins). The binding between MDGA2 and GluN1 has not been previously reported, and therefore wondered if this may be a direct interaction. We found that MDGA2 and GluN1 can directly interact in heterologous cells and confirmed that MDGA2 binds to GluN1 with higher affinity than MDGA1 (Fig. 3C), mirroring the finding that GluN1 is enriched in the MDGA2, but not the MDGA1, proteome. Next, we tested whether NMDAR subunit composition affects the preference of binding to MDGA2 *vs* MDGA1 and found that both GluN1/GluN2A (Fig. 3D) and GluN1/GluN2B (Fig. 3E) complexes interact preferentially with MDGA2 over MDGA1, with C, co-IP of overexpressed Flag-tagged GluN1 subunit of the NMDAR with HA-tagged MDGA1/2 in HEK293T cells. Quantification is shown to the right. Interaction with MDGA2 is of approximately 3-fold higher affinity than MDGA1 (p<0.001). D, co-IP of overexpressed Flag-tagged GluN2A subunit of the NMDAR together with the obligatory GluN1 subunit with HA-tagged MDGA1/2 in HEK293T cells. Quantification is shown to the right. Both GluN2A and GluN1 interact with MDGA2 with higher affinity than with MDGA1 (p=0.0037 andp=0.0092, respectively). E, co-IP of overexpressed Flag-tagged GluN2B subunit of the NMDAR together with the obligatory GluN1 subunit the with HA-tagged MDGA1/2 in heterologous HEK cells. Quantification is shown to the right. Both GluN2B and GluN1 interact with MDGA2 with higher affinity than with MDGA1 (p<0.0001 and p=0.0105, respectively). n=3 for all experiments. *, p<0.05; **, p<0.01; ***, p<0.001 (Student’s T-test). “WB”, Western blot; “Ab”, antibody.

**Fig 3.**
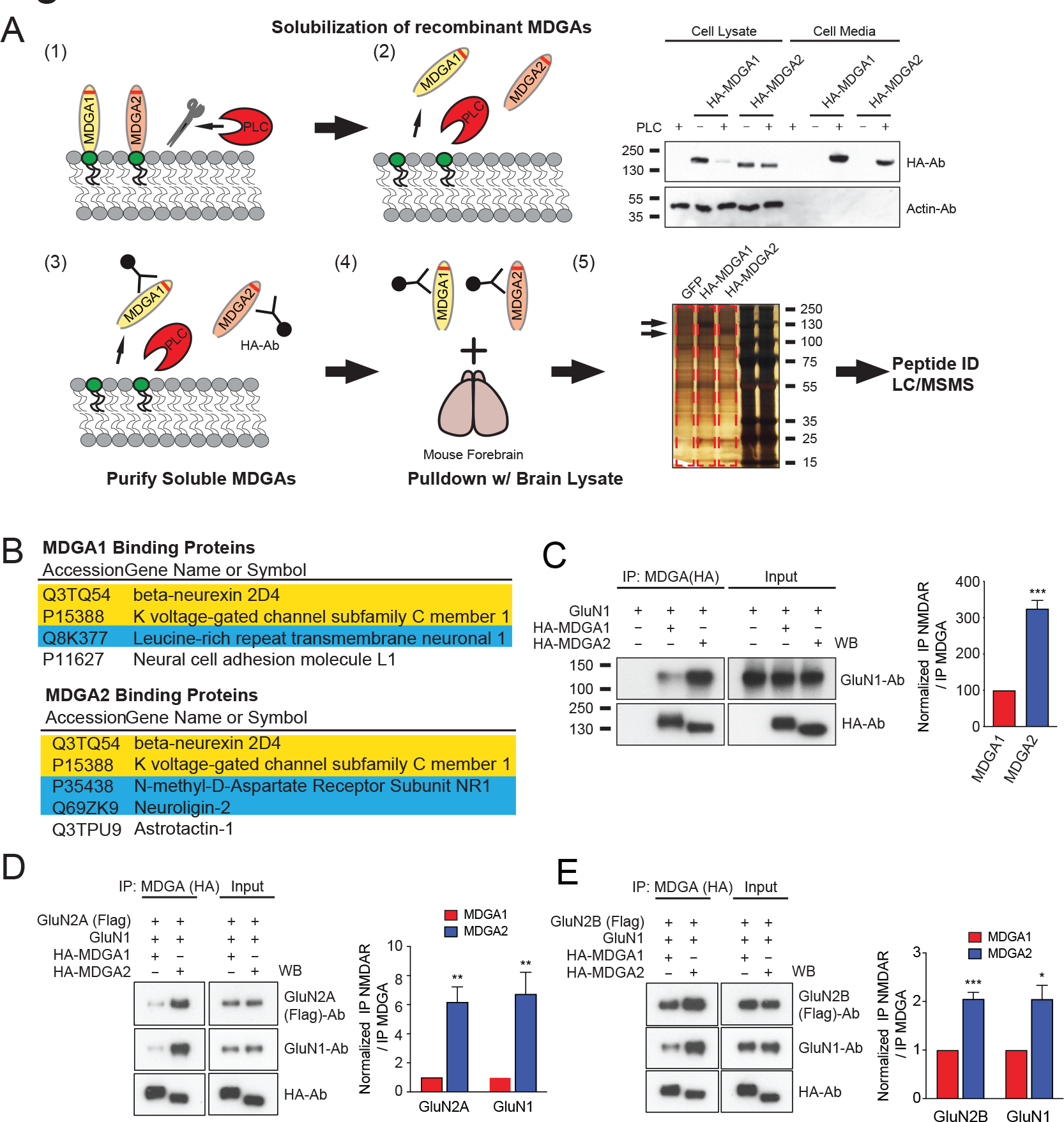
**Unbiased screen to identify MDGA1 and MDGA2 interacting proteins identifies isoform-specific interactions between MDGA proteins and NMDAR subunits**. A, Five-step process to identify MDGA interacting proteins. (1) MDGA1 or MDGA2 were expressed in HEK293T cells and treated with phospholipase C (PLC) to release the MDGAs from the membrane. (2) Immunoblot analysis of transfected HA-MDGAs ± PLC showing a reduction in cell lysate and an increase in cell media of MDGAs following PLC treatment. (3) MDGAs were precipitated from the Media with an HA-Ab. (4) Purified MDGAs were incubated with solubilized brain lysate. (5) Interacting proteins were visualized with silver staining and identified using mass spectrometry. Arrows indicate MDGA1 and MDGA2 respectively. B, List of representative MDGA1 and MDGA2 interacting proteins that were pulled down in at least 2 of 3 experiments (see Supplementary Table 1 for the complete list). Yellow highlight indicates association with both MDGA1 and MDGA2. Blue highlight indicates proteins exclusively found to interact with MDGA2 either found previously in the literature or confirmed with co-IP. Bar graphs plot transfected amplitude normalized to control cell ± SEM. E, AMPAR (p=0.0010, n=13)- and NMDAR (p>0.05, n=12)-mediated EPSC scatter plots displaying a selective reduction in only AMPAR-mediated EPSC amplitudes in MDGA1 KD transfected cells compared to control cells. F, AMPAR (p>0.05, n=11)- and NMDAR (p=0.008, n=10)- mediated EPSC scatter plots displaying a selective reduction in only NMDAR-mediated EPSC amplitudes in MDGA2 KD transfected cells compared to control cells. **, p<0.01; ***, p<0.001, Wilcoxon signed-rank test. Scale bar for D-F: 25pA, 0.1s. “WB”, Western blot; “Ab”, antibody; “DG”, dentate gyrus.

GluN1/GluN2A receptors showing stronger preference. Lastly, we assessed whether the GluN1 subunit is necessary for MDGA2-NMDAR interaction and found that both GluN2A and GluN2B can interact with MDGA2 independently of GluN1 (Fig. S4). Collectively, these data support MDGA2 localization at both excitatory and inhibitory synapses.

### The role of MDGAs in excitatory synaptic transmission

Our present results, together with those of others, indicate that MDGAs localize, at least in part, to synapses. However, their role in synaptic transmission is controversial. We therefore examined the physiological consequences of deleting MDGA1 and MDGA2 either individually or together (to control for the potential for functional redundancy due to their high sequence homology) on excitatory and inhibitory synaptic transmission. CA1 PNs express MDGAs [Figs. 1 and 2, (*14, 20*)], thereby constituting an ideal cell type to elucidate their cell-autonomous role in synaptic transmission.

We initially confirmed that the MDGA shRNAs (*13*) selectively reduce MDGA protein (Fig. 4A). To explore the physiological role of MDGA proteins on excitatory synaptic transmission, we biolistically transfected shRNAs targeting MDGA1 and MDGA2 into organotypic hippocampal slice cultures, which results in sparse transfection of a few CA1 PNs per slice. AMPA receptor (AMPAR)- and NMDAR-mediated EPSCs, evoked with a stimulating electrode placed in SR, were simultaneously measured in a KD cell and a neighboring, non-transfected control CA1 PN was held at -70 mV and +40 mV, respectively (Fig. 4B-C). The simultaneous KD of MDGA1 and MDGA2 decreased both AMPAR and NMDAR mediatedcurrents by approximately 50% (Fig. 4D). To assess whether MDGA1 or MDGA2 alone accounted for these effects, we performed individual KD of each protein. MDGA1 KD alone wassufficient to recapitulate the impairment in AMPAR EPSCs caused by the dual deletion, while leaving NMDAR EPSCs unaffected (Fig. 4E). Conversely, MDGA2 KD reproduced the decrease in NMDAR EPSCs found in the dual KD, while leaving AMPAR currents unaltered (Fig. 4F).

**Fig 4.**
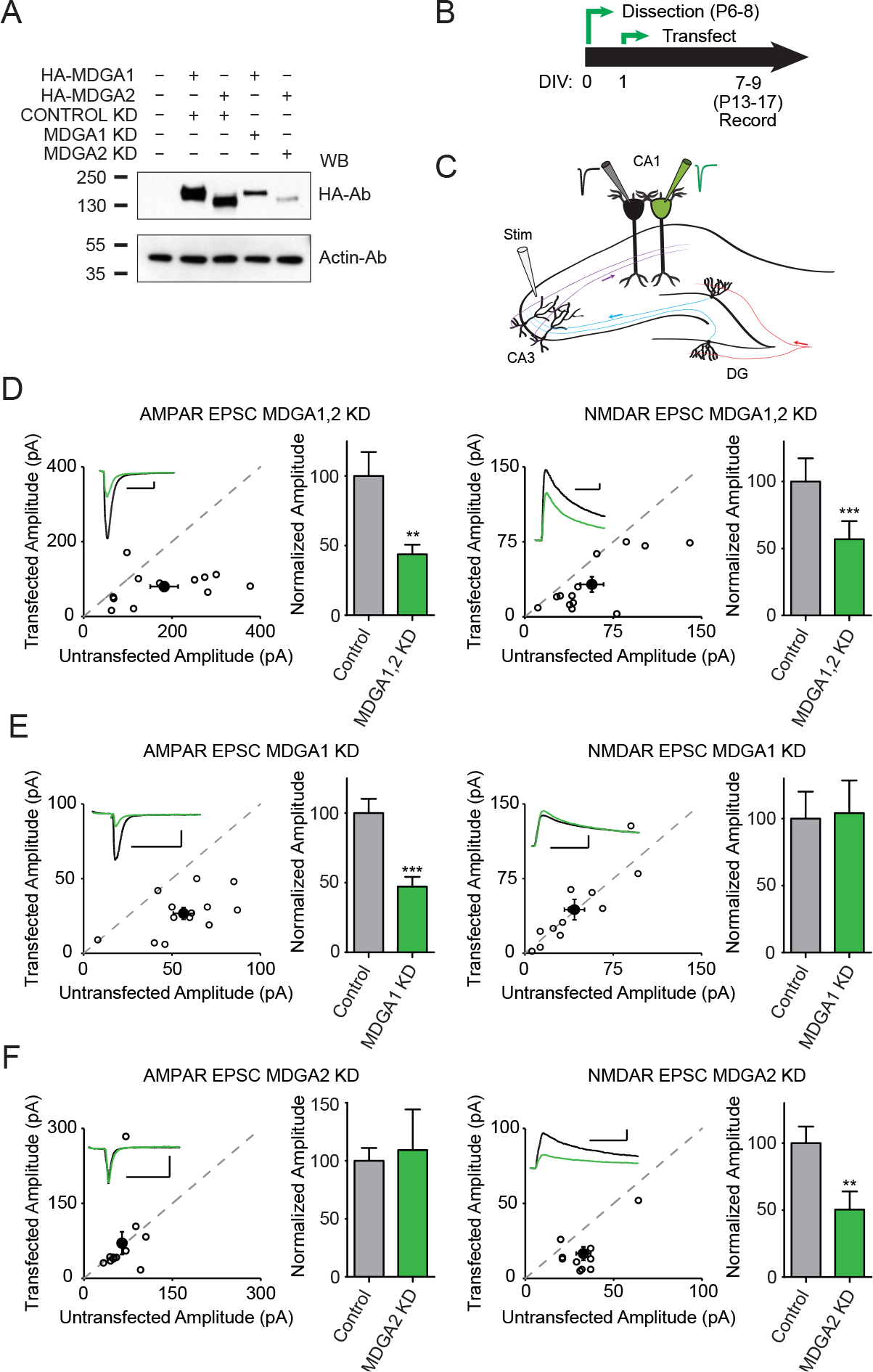
**Knockdown of MDGA family decreases excitatory currents.** A, Immunoblot analysis of HA-MDGAs transfected with MDGA1, MDGA2, or control shRNA in HEK cells. B, Experimental timeline. C, Dual whole-cell recording setup in organotypic hippocampal rat slices. Black and green filled neurons represent untransfected (control) and transfected (experimental) neurons, respectively. D, AMPAR (p, =0.0049, n=12)- and NMDAR (p=0.0005, n=13)-mediated EPSC scatter plots displaying reductions in MDGA1,2 KD transfected cells compared to control cells. Open circles are individual pairs, filled circle is mean ± SEM. Representative traces show control (black) and transfected (green) neurons.

These findings indicate that MDGA1 and MDGA2 are essential for excitatory synaptic transmission in the SC-CA1 PN synapse and selectively modulate AMPAR- and NMDAR-mediated currents, respectively. Strikingly, these results are the opposite of what one would expect if MDGAs are synaptic repressors.

### Role of MDGAs in inhibitory synaptic transmission

We explored the effect of combined KD of MDGA1 and MDGA2 in inhibitory synaptic currents (Fig. 5A). Cells lacking both MDGA1 and MDGA2 exhibited enhanced inhibitory postsynaptic currents (IPSCs, Fig. 5D), with the individual KD of MDGA2 (Fig. 5F), but not MDGA1 (Fig. 5E), selectively increasing inhibitory currents. These findings are consistent with a role for MDGAs as synaptic repressors at inhibitory synapses, yet this effect is confined to MDGA2, and not to MDGA1 as previously suggested (*12, 17, 18, 20*). Having identified a synaptic repressor role for MDGA2, we tested whether repressor function is mediated via the interaction with Nlgn2 which we identified using unbiased proteomics (Fig. 3B). To do so, we mutated the phenylalanine and tyrosine amino acids in positions 149 and 150 of MDGA2 with alanine (FY149-150AA), thereby blocking the previously found interaction with Nlgn2 (*24*). We first demonstrated that expressing a shRNA-resistant version of MDGA2 (Fig. 5B, C) fully reverses the enhancement of IPSCs seen in the KD (Fig. 5H). Conversely, expression of the FY149-150AA mutant together with the MDGA2 shRNA resulted in a robust enhancement of IPSCs (Fig. 5I) underscoring that the repressor function is dependent on the neuroligin interaction.

**Fig 5.**
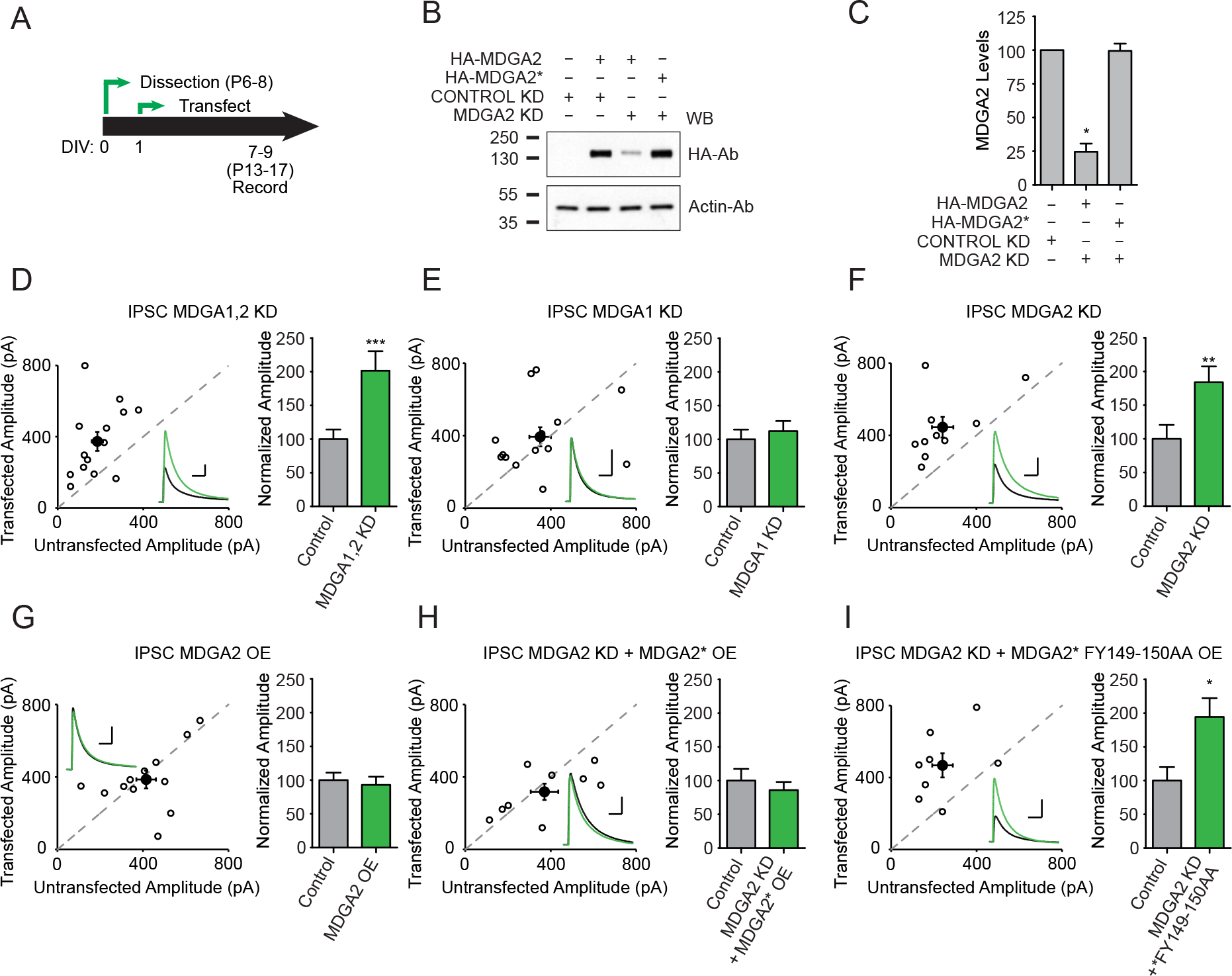
**Knockdown of MDGA2 increases inhibitory currents in a neuroligin-dependent manner**. A, Experimental timeline. B, Immunoblot analysis of HA-MDGA2 or HA- MDGA2* (shRNA resistant plasmid) transfected with MDGA2 or control shRNA in HEK cells. C, Total MDGA2 lysate levels (means ± s.e.m.) normalized to control show efficient MDGA2 KD against HA-MDGA2 (p=0.0286, n=4), but not HA-MDGA2* (p>0.9999). D, Scatter plots showing enhancements in IPSC-mediated currents in MDGA1,2 KD transfected cells compared to untransfected control cells (p=0.0009, n=14). Open circles are individual pairs, filled circle is mean ± s.e.m. Black sample traces are control, green are transfected neurons. Bar graphs plots transfected amplitude normalized to control cell ± s.e.m. E, Scatter plots showing no significant difference in IPSC-mediated currents in MDGA1 KD transfected cells compared to untransfected control cells (p>0.05, n=14). F, Scatter plots showing enhancements in IPSC-mediated currents in MDGA2 KD transfected cells compared to untransfected control cells (p=0.002, n=10). G, Scatter plots showing no significant difference in IPSC-mediated currents in MDGA2 overexpressing (OE) transfected cells compared to untransfected control cells (p>0.05, n=12). H, Scatter plots showing no significant difference in IPSC-mediated currents in MDGA2 KD + MDGA2* OE transfected cells compared to untransfected control cells (p>0.05, n=9). I, Scatter plots showing enhancements in IPSC-mediated currents in MDGA2 KD + MDGA2* FY149-150AA OE transfected cells compared to untransfected control cells (p=0.0391, n=9). IPSC amplitudes recorded at 0 mV. *, p<0.05; **, p<0.01; ***, p<0.001, Mann-Whitney U test (C), Wilcoxon signed-rank test (D-I). Scale bar for D-I: 100 pA and 0.05 s. “WB”, Western blot. “Ab”, antibody.

### Role of MDGAs in dendritic spine morphology

Anatomically, co-expression of the shRNAs for MDGA1 and MDGA2 did not change spine density in CA1 PNs nor did it alter the spine head diameter (Fig. S5A-D), although it caused a significant reduction in neck length and an increase in neck diameter (Fig. S5E, F). Thus, MDGA proteins appear to play a role in regulating spine neck morphology, thereby potentially regulating biochemical and electrical spine compartmentalization and signaling (*28, 29*).

### Synaptic effects of overexpressing MDGAs

Our results are provocative because they appear to contradict previous findings, many of which were obtained using overexpression of MDGAs. First, our data show a dramatic functional specificity of MDGA1 and MDGA2 at excitatory and inhibitory synapses and, second, while the action of MDGA2 at inhibitory synapses confirms the synaptic repressor role, the action of MDGA1 at excitatory synapse is fundamentally different. To resolve these apparent contradictions, we carried out experiments using overexpression of MDGAs. Remarkably, overexpression of MDGA1 (Fig. S6A, B) *reduced* both AMPAR and NMDAR EPSCs (Fig.S6C), thus indicating that overexpressed MDGAs behave as synaptic repressors, as expected from previous reports. A limitation of overexpression experiments is that high levels may cause protein mistargeting and ultimately “gain-of-function” for the examined protein, a reasonable possibility for MDGAs given the difference seen in endogenous *vs.* exogenous localization studies [see figures above, (*12, 13, 15, 17*)]. To address this concern, we inserted an internal ribosomal entry site (IRES) sequence upstream of the MDGAs cDNA (Fig. S6A-B), a strategy which limits protein expression (*30, 31*). Consistent with previous findings (*17*), but in contrast with MDGA2 overexpression (Fig. 5G), we found that both strong (Fig. S7A) and mild (Fig. S7B) MDGA1 overexpression causes a marked reduction of IPSCs. These findings are consistent with MDGA1 acting as a synaptic repressor at inhibitory synapses. Given previous reports suggesting that MDGA1 function requires interaction with Nlgn2 (*12, 17*), we tested whether an interaction between MDGA1 and Nlgn2 was required for the MDGA1 overexpression-induced reduction in synaptic transmission. Interestingly, we found that overexpressing the MDGA1 FY147-148AA mutant that lacks the Nlgn2-binding motif results in AMPAR and NMDAR EPSCs being as strongly decreased as with WT MDGA1 overexpression (Fig. S6E), while it no longer affects IPSCs (Fig. S7C). These findings indicate that neuroligin interaction is required for the depressive effect of MDGA1 overexpression, yet only at inhibitory synapses.

### CRISPR/Cas9 deletion of MDGAs

Our findings indicate that both endogenous MDGAs support excitatory synaptic transmission and that MDGA2 acts as a repressor of inhibitory synaptic transmission. However, manipulations involving shRNAs can have off-target effects (*32*). Therefore, to verify our shRNA results we used an alternative genetic deletion approach, based on sparse CRISPR/Cas9- mediated KO in organotypic hippocampal slices (*33*). We designed several gRNAs against the MDGAs, tested their KO efficiency, and selected the best for further evaluation (Fig. 6A, B).

**Fig 6.**
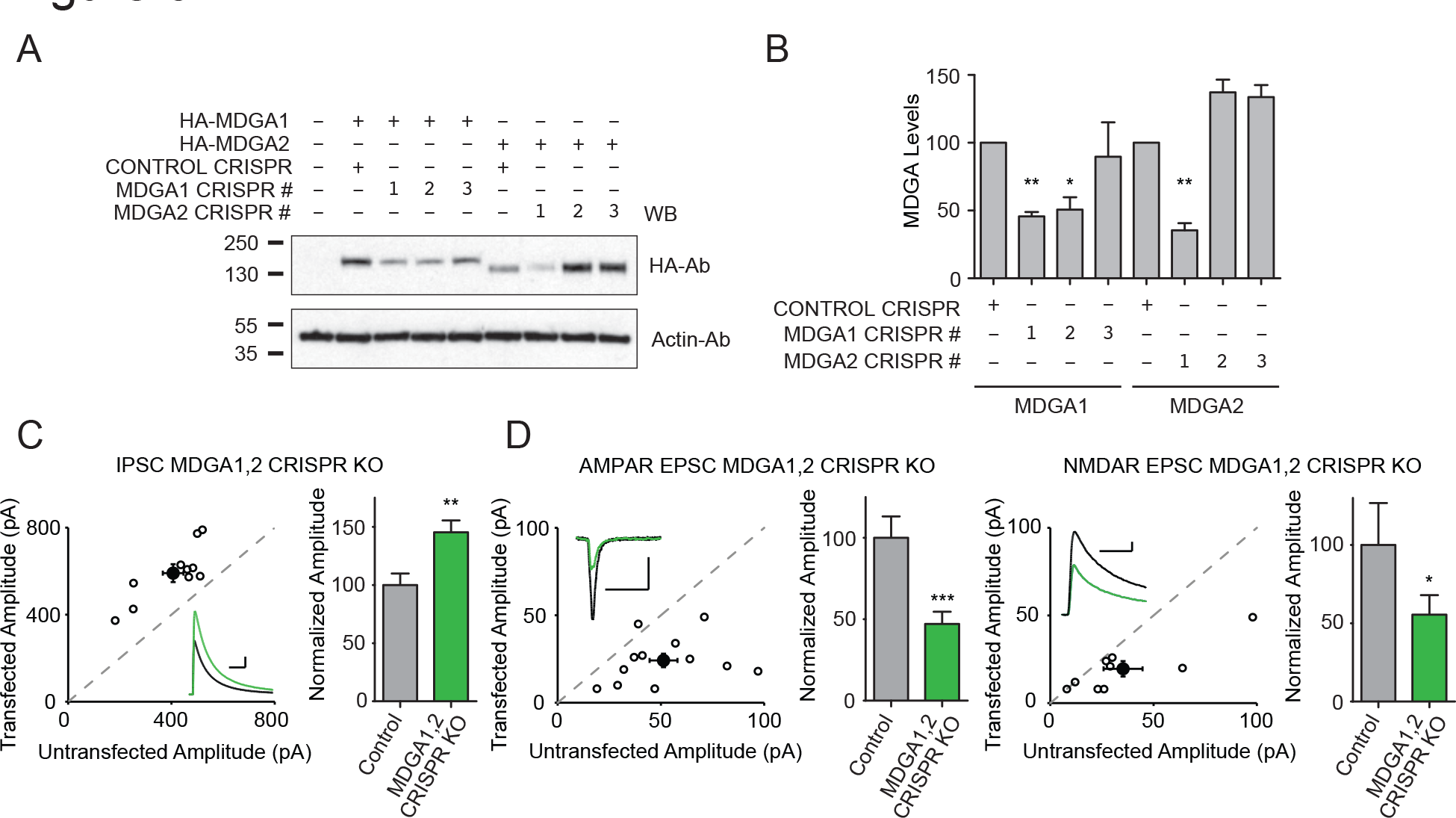
**CRISPR knockout of MDGA family increases inhibitory and decreases excitatory currents**. A, Immunoblot analysis of HA-MDGAs transfected with MDGA1, MDGA2, or control CRISPRs in HEK cells. B, Total MDGA lysate levels (means ± SEM) normalized to control show efficiency of MDGA1 CRISPR #1 (p=0.0035, n=5) and MDGA1 CRISPR #2 (p=0.0328, n=5), and MDGA2 CRISPR #1 (p=0.0065, n=5). C, Scatter plots showing enhancements in IPSC-mediated currents in MDGA1,2 KO transfected cells compared to untransfected control cells (p=0.0020, n=10). Open circles are individual pairs, filled circle is mean ± SEM. Black sample traces are control, green are transfected neurons. Scale bar denotes 100 pA and 0.05 s. Bar graphs plots transfected amplitude normalized to control cell ± SEM. D, AMPAR (p=0.0010, n=12)- and NMDAR (p=0.0156, n=9)-mediated EPSC scatter plots displaying reductions in MDGA1,2 KO transfected cells compared to control cells. Scale bar denotes 25 pA and 0.1 s. Bar graphs as in C. *, p<0.05; **, p<0.01; ***, p<0.001, Mann-Whitney U test (B), Wilcoxon signed-rank test (C-E). “WB”, Western blot; “Ab”, antibody.

Using an analogous experimental strategy as the one utilized in our shRNA KD experiments, we analyzed the effect of the dual MDGA1/2 CRISPR on synaptic transmission. This alternative method resulted in an increase in IPSCs (Fig. 6C), and a reduction in AMPAR- and NMDAR-EPSCs (Fig. 6D). These results are indistinguishable from those obtained with the shRNA approach, thus solidifying our conclusion that MDGAs act as synaptic repressors only at inhibitory synapses and are facilitators of synaptic transmission at excitatory synapses.

## Discussion

The formation, maintenance, and activity-dependent modification of synapses is orchestrated by a multitude of synaptic adhesion molecules, commonly referred to as “synaptic organizers” (*4*). Broadly, synaptic organizers perform two functions: inducing the assembly of new synapses or specifying synapse properties (*27*). The recent proposal that MDGAs act as synapse repressors suggests an additional, perhaps underappreciated layer of synapse regulation. While there are multiple adhesion molecules that have been demonstrated to positively influence synapse formation, MDGAs are receiving substantial attention as negative regulators, a role only few other synaptic proteins have been shown to play (*34–36*). Elucidating the endogenous functions of MDGAs has not been a trivial task, as simply localizing MDGAs to the synapse has not been straightforward.

Ten years after the first reports of the synaptic roles of MDGAs (*12, 17*), the precise cellular localization and role of endogenous MDGAs remains enigmatic. Similar to other synaptic cell adhesion molecules, the function(s) of MDGAs were initially defined largely based on overexpression studies (*12, 13, 17*) although, taken together, these studies underscore the difficulties in interpreting overexpression experiments. Perhaps this is particularly the case for MDGAs because of their lack of intracellular domains allowing direct anchoring to excitatory (i.e. through MAGUKs) or inhibitory (i.e. through gephyrin/collybistin) synapses, making them more prone to mislocalization when overexpressed. *In situ* hybridization and β-galactosidase reporter experiments confirm that the MDGA1 and MDGA2 are expressed in neurons (*11, 17, 37*) (*14, 19, 20*). Proximity-based proteomic assessment of endogenous MDGA localization found that MDGA1 was localized to excitatory synapses, while MDGA2 was localized to inhibitory synapses in cultured neurons (*13*). Additionally, KD strategies have been used to gain insight on the physiological role of MDGA proteins. Several reports (*12, 13, 17*) found increases in inhibitory synapse density in dissociated neuronal cultures after MDGA1 KD, establishing the prevailing view that MDGA1 acts primarily to negatively regulate inhibitory synapses. In contrast, more recent KO studies challenge this notion reporting that i) deletion of MDGA1 does not alter inhibitory synapse number or transmission (*18*) and ii) loss of either MDGA isoform did not affect IPSCs (*15*). In sum, although there is general consensus in the field that MDGAs can act as synaptic repressors, these seemingly divergent sets of results constrain the formation of an integrative model of MDGA synaptic function.

Therefore, when we set out to elucidate their function, it was imperative to establish their subcellular localization. We generated epitope-tagged MDGA1 and MDGA2 KI mice to circumvent technical limitations with MDGA antibodies. Our results represent a breakthrough in the study of endogenous MDGA2, by allowing the detection of the endogenous protein by immunoblot. We uncovered that both MDGA proteins are widely expressed throughout the CNS and follow a very marked developmentally regulated expression pattern. Specifically, the chronological and regional expression pattern of MDGA1 and MDGA2 are largely overlapping (noteworthy given they do not appear to have overlapping synaptic functions), with a peak of expression around the 2^nd^-3^rd^ weeks of postnatal life. Remarkably, our imaging data provide the first direct evidence of the enrichment of MDGA1 at SC-CA1 PN synapses in the hippocampus and indicates that endogenous MDGA1 is expressed, at least in part, at young excitatory synapses *in vivo*. Intriguingly, we identify the existence of a significant pool of MDGA1 which is not at excitatory or inhibitory synapses, as previously suggested (*15*). This potentially non-synaptic pool of MDGA1 is deserving of future exploration. Disappointingly, our genetic approach did not allow for imaging of MDGA2 in intact brain tissue. However, our proteomics, biochemistry, and electrophysiology data are consistent with previous proteomic analyses of endogenous MDGA2 localization (*13*), and deem it likely that MDGA2 is present at both excitatory and inhibitory synapses onto hippocampal CA1 PNs. Interestingly, few postsynaptic cell adhesion molecules are shared by excitatory and inhibitory synapses. To the best of our knowledge, neuroligin-3 (Nlgn3) is the only other exception to the rule (*38*). Unlike Nlgn3, MDGA2 plays fundamentally different roles at each synapse type, acting as a repressor at inhibitory synapses but supporting NMDAR transmission at excitatory synapses, again highlighting the uniqueness of the synaptic roles played by MDGA2.

The disparate results seen for MDGA KD between neuronal cultures and *in vivo* experiments coupled with emerging evidence that synaptic organizers can have different roles at different synapse types, we performed all the physiology experiments on a well-defined preparation and synapse type (SCèCA1 PN synapse in mouse hippocampal slices) to mitigate these confounding factors. We found that the individual KD of MDGA1 and MDGA2 cell-autonomously and selectively decreases AMPAR- and NMDAR-mediated currents, respectively. We are not aware of another family of synaptic organizers with such diverse roles at the same synapse. Conversely, eliminating MDGAs from neurons enhanced IPSCs, an effect entirely mediated by MDGA2 deletion, supporting our proposal that endogenous MDGA2 acts as a repressor of IPSCs. Together with our imaging, proteomics and biochemistry data, these results provide converging evidence for the localization of MDGA1 at excitatory synapses, and of MDGA2 at both excitatory and inhibitory synapses. These findings are largely consistent with the endogenous localization study of Loh *et al.* (*13*), and with the notion that MDGAs act as repressors at inhibitory synapses (*12, 16, 17*). Provocatively, these data show that, in contrast to what had previously been suggested (*14, 15*), MDGAs do not act as repressors of excitatory synapses and have a fundamentally different role in maintaining EPSCs.

Albeit through a fundamentally different mechanism to the previously proposed, our results are largely consistent with findings using MDGA KO mice. For example, the decreased E/I ratio found in MDGA1 KO mice can be explained by an increase of inhibitory synapses, as suggested previously (*20*) but also by a decrease in excitatory transmission (present study). The altered LTP and learning in MDGA1 KOs can be explained by the reduced AMPAR trafficking or function found in our study. Similarly, the reported increase in AMPAR/NMDAR ratio in the MDGA2 heterozygotes (the homozygous KO is lethal), previously associated with an increase in AMPAR levels (*14*) is also consistent with a specific decrease in NMDAR EPSCs reported herein. The impaired NMDAR function could conceivably also influence the deficits in LTP and hippocampal-dependent learning and memory found in the MDGA2 heterozygotes (*14*).

Interestingly, our KD studies revealed a previously unrecognized role of MDGA proteins regulating dendritic spine morphology, which may also contribute to the MDGA KO mice phenotype. Specifically, MDGA1/2-lacking spines have significantly shorter and wider spine necks. Alterations in spine morphology are linked with numerous neurological disorders including schizophrenia and ASD, two disorders with which MDGA1 and MDGA2 are associated, respectively (*39*).

Previous reports indicate that MDGA1’s function depends on the modulation of neuroligin-neurexin interactions (*12, 15, 17, 22-24*). However, our study failed to find a functional relationship between endogenous MDGA1 and Nlgn2. Specifically, we did not find i) an effect of MDGA1 KD in IPSCs, ii) a co-distribution of MDGA1 at Nlgn2-positive puncta in the mouse brain, and iii) Nlgn2 enriched in the unbiased MDGA1 proteome. This is consistent with a recent report which suggested that, rather than neuroliginss, MDGA1 function requires its interaction with presynaptic amyloid precursor protein [APP, (*18*)]; although, to note, APP was not found in our immunoprecipitation studies (Supplementary Table 1). We did find that exogenous MDGA1 expression leads to decreased IPSCs, in accord with previous reports (*12, 17, 22*). However, these overexpression data, including our own, are inconsistent with MDGA1 KD/KO data in which deleting MDGA1 has no effect on synaptic inhibition [present study, (*15, 18*)]. Therefore, we conclude that overexpression approaches have led to substantial confusion about synaptic MDGA localization, perhaps through “gain of function” effects by mistargeting of the protein to non-endogenous synapses, a possibility consistent with our finding that the repressive effect of MDGA1 overexpression at inhibitory synapses is abrogated by mutations that diminish binding with Nlgn2, a protein endogenous MDGA1 likely does not share spatial proximity with but with which it may interact when overexpressed (Fig. S7).

Instead, we found that, in addition to NMDAR-mediated EPSCs, MDGA2 controls inhibitory synaptic transmission. Interestingly, a MDGA2 mutant with impaired interaction with Nlgn2 did not rescue IPSCs in a MDGA2 KD background, in contrast with the full rescue achieved with WT MDGA2. This finding is consistent with the MDGA2-Nlgn2 interaction revealed in our unbiased proteomics. Altogether, our results challenge the model where MDGA1 acts as a specific gatekeeper of inhibitory synapse formation and/or function (*20, 21, 25*) and are consistent with a more prominent role at excitatory synapses. Our findings instead indicate that the gatekeeper of inhibitory synapses is MDGA2, which likely involves the regulation of Nlgn2. Furthermore, we uncovered that MDGA2 is required for NMDAR function. We found that both the obligatory GluN1 subunit and the GluN2A and GluN2B subunits can directly bind MDGA proteins independently, and that they show a preference for MDGA2 *vs* MDGA1 in a co-immunoprecipitation assay, again an intriguing result given their high sequence conservation.

However, the finding that only GluN1 is represented in the MDGA2 proteome from native mouse brain tissue indicates that this subunit may play the predominant role in the MDGA2- NMDAR interaction *in vivo*. This raises an interesting scenario in which two different synaptic MDGA2 pools coexist, one specialized in regulating NMDAR (neuroligin-independent) and another dedicated to modulate GABAergic transmission (neuroligin-dependent).

How do MDGAs contribute to excitatory synaptic transmission? The elegant structural studies on the MDGA/neuroligin complex (*22–24*) provide a molecular explanation for our finding that MDGA2 acts as a synaptic repressor at inhibitory synapses. In contrast, the mechanisms underlying the action of MDGAs at excitatory synapses remain uncertain. As described above, MDGAs lack transmembrane or intracellular domains, which typically direct synaptic targeting. Therefore, they are likely to rely on extracellular protein-protein interactions for their synaptic localization, which presumably underlies their relatively low synaptic localization compared to other synaptic proteins [our data, (*15*)]. The interaction between MDGA2 and NMDAR thus conceivably involves extracellular NMDAR motifs, which are critical for receptor trafficking and function (*40*). Similarly, recent work has established a prominent role for extracellular domains for synaptic AMPAR localization (*41–43*). These findings suggest that extracellular interactions with synaptic cleft proteins, including synaptic adhesion molecules such as MDGA1, may regulate AMPAR trafficking and function (*6, 44*).

Although we did not find a direct interaction between AMPARs and MDGA1, we did find that LRRTM1 was found specifically in the MDGA1 proteome. This is noteworthy for several reasons. First, this protein specifically promotes AMPAR transmission without affecting NMDAR function (*45*), mimicking MDGA1’s role. Second, the LRRTM proteins were used as the “bait” which unbiasedly and indirectly localized MDGA1 at excitatory synapses (*13*). Third, as with MDGA1, LRRTM1 is associated with schizophrenia (*46*). Collectively, the relationship between LRRTMs and MDGAs remains an exciting area of future exploration. Notably, even after MDGAs bind to neuroligins via their Ig1 and Ig2 domains, they still retain large interacting surfaces and are likely poised for other potential molecular interactions. The most intriguing candidates from our proteomics work suggest those may happen in *cis* like neuroligins (GluN1, LRRTM1); however, our unbiased approach also pulled out synaptic proteins like neurexins which would presumably interact in *trans* across the synapse, an exciting avenue of future exploration.

Summarily, our study provides the first subcellular localization data of endogenous MDGA1 in brain tissue and establishes that MDGA proteins play essential, yet highly divergent roles at different synapse types. Future directions will include learning how MDGA1 and MDGA2 control AMPAR and NMDAR transmission, as well as explore the role of MDGA proteins in other brain circuits as identified by the expression of MDGAs found in our KI mice. The overall theme to emerge from our work is that MDGAs do not perform a unitary function (i.e., repressors) at synapses. Instead, the different MDGA family members play unique and complex roles in shaping transmission at excitatory and inhibitory synapses.

## Materials and Methods

### Animals

All animal procedures were approved by the Institutional Animal Care and Use Committees at the University of California, San Francisco (protocol number AN183289, PI, Roger Nicoll) and University of California, Irvine (protocol number AUP-20-156, PI, Javier Diaz-Alonso). All animals were maintained in 12 hour (h) light/dark schedule and with access to food and water, *ad libitum*.

Generation of HA-MDGA1 and Myc-MDGA2 mouse strains was performed by Cyagen Inc. by CRISPR/Cas9 mediated homology-directed repair (Fig. S1A and Fig. S2A). Briefly, the gRNA to mouse Mdga1 gene (5’-CCCTTCCACTGTCGGGGACAAGG-3’), the donor oligo containing the HA-tag-RSRD linker (5’-TACCCATACGATGTTCCAGATTACGCTAGATCTCGAGAT-3’) flanked by 120 nt homology arms combined on both sides and Cas9 mRNA were coinjected into fertilized mouse eggs to generate targeted knock-in (KI) offspring. F0 founder animals were identified by PCR followed by sequence analysis, and bred to WT mice to test germline transmission and F1 animal generation. At least 5 backcrossings were performed before using the animals for experiments to minimize the possible artifacts caused by non-specific insertions. An analogous procedure was followed for Myc-MDGA2, using the gRNA 5’ TCCACTCACCGTACACTCCTTGG-3’ and the Myc-tagged-RSRD linker 5’- GAACAAAAACTCATCTCAGAAGAGGATCTGAGATCTCGAGAT-3’.

Validation of the successful KI was achieved by genotyping PCR (performed by TransnetYX, Inc.), genomic sequencing (Figs. S1B, S2B), Western blot, and immunofluorescence. The predicted protein sequences after successful recombination are indicated in Fig. S1C and Fig. S2C, respectively. After backcrossing, both colonies were maintained in homozygosity and a HA-MDGA1/Myc-MDGA2 colony was created and used for most of the experiments in this paper. Postnatal day (P) 3-130 HA-MDGA1/Myc-MDGA2 mice of either sex were used in this study.

P6-8 (Sprague Dawley) rat pups of either sex were employed to generate the organotypic hippocampal slice cultures employed in MDGA overexpression, shRNA-mediated MDGA KD and CRISPR/Cas9-mediated MDGA KO experiments.

### Constructs

Rat pCAG-HA-MDGA1, Mouse pCAG-HA-MDGA2 (generous gift from Ann Marie Craig’s Laboratory, University of British Columbia), pCAG-HA-MDGA2* (shRNA resistant), pCAG-HA-MDGA1 FY147-148AA, pCAG-HA-MDGA2* FY149-150AA, pCAG-mCherry-IRES-HA-MDGA1, pCAG-mCherry, GluN1-GFP (generous gift from Stephen Traynelis’s laboratory, Emory University) were used for biochemical and electrophysiology experiments. The primers used to create MDGA2* were forward 5’- AGTATAGGCGAGGCCAAGGAGCAGTTTTAC - 3’ and reverse 5’- GTAAAACTGCTCCTTGGCCTCGCCTATACT- 3’, MDGA1 FY147-148AA were forward 5’- GCGACGTCCGAGGCAACGCCGCCCAGGAGAAGACCGTGT - 3’ and reverse 5’- ACACGGTCTTCTCCTGGGCGGCGTTGCCTCGGACGTCGC- 3’, MDGA2 FY149-150AA were forward 5’- TATAGGCGAGGCCAAGGAGCAGGCTGC CTATGAGAGAACAGTGTTCCTC - 3’ and reverse 5’- GAGGAACACTGTTCTCTCA TAGGCAGCCTGCTCCTTGGCCTCGCCTATA- 3’, IRES-HA-MDGA1 were forward 5’-CTTGCCACAACCCGGGATGGATGTCTCTCTTTGCCC - 3’ and reverse 5’- CTCGAGCTAGCGGCCGCTCATCTCTGCAACGCCAAGA- 3’, IRES-HA-MDGA2 were forward 5’- CTTGCCACAACCCGGGATGGATGTCTCTCTTTGCCC - 3’ and reverse 5’- CTCGAGCTAGCGGCCGCTCACCTTCGAGGGCTTAAGA- 3’. pLLS-anti MDGA1 and pLLs-anti MDGA2, which dually express GFP for positive transfection identification (generous gifts from Alice Ting’s Laboratory, Stanford University), knockdown (KD) constructs were used for electrophysiology and imaging. Knockout (KO) constructs used for electrophysiology and biochemistry were MDGA1 KO CRISPR #1 (sequence: TCCGGGAGAGCGACACCCTG) MDGA1 KO CRISPR #2 (sequence: GACGGTACAGCGTAGAAACA), MDGA1 KO CRISPR #3 (sequence: GATAAAGCGGGCGGGCGGGT), MDGA2 KO CRISPR #1 (sequence: AGCAATAAAGTCGATCCGAG), MDGA2 KO CRISPR #2 (sequence: ACTCGGATCGACTTTATTGC), and MDGA2 KO CRISPR #3 (sequence: TACAGTAATATCGGCCTCCT). The CRISPR constructs were generated using a standard PCR cloning procedure that included fusing antisense primers and subcloning into a lentiviral vector that expressed Cas9 and GFP. The primers used to create MDGA1 KO CRISPR #1 were forward 5’- CACCGCAGGGTGTCGCTCTCCCGGA – 3’ and reverse 5’- AAACTCCGGGAGAGCGACACCCTGC – 3’, MDGA1 KO CRISPR #2 were forward 5’ – CACCGACGGTACAGCGTAGAAACA – 3’ and reverse 5’- AAACTGTTTCTACGCTGTACCGTC – 3’, MDGA1 KO CRISPR #3 were forward 5’ - CACCGATAAAGCGGGCGGGCGGGT – 3’ and reverse 5’- AAACACCCGCCCGCCCGCTTTATC – 3’, MDGA2 KO CRISPR #1 were forward 5’- CACCGAGCAATAAAGTCGATCCGAG – 3’ and reverse 5’- AAACCTCGGATCGAC TTTATTGCTC – 3’, MDGA2 KO CRISPR #2 were forward 5’ - CACCGACTCGGAT CGACTTTATTGC – 3’ and reverse 5’- AAACGCAATAAAGTCGATCCGAGTC – 3’, MDGA2 KO CRISPR #3 were forward 5’ – CACCGTACAGTAATATCGGCCTCCT – 3’ and reverse 5’- AAACAGGAGGCCGATATTACTGTAC- 3.

### Cell culture and transfections

HEK293T cells (ATCC) were grown and maintained in DMEM (Gibco, 11966025) supplemented with 10% fetal bovine serum [FBS (Hyclone, SH30071.03)] and 1% glutamine (Gibco, 25030081) without antibiotic in a humidified incubator at 37°C with 5% CO2. For biochemistry, transfections were performed directly after splitting the cells in 6-well plates using Lipofectamine 2000 Reagent (Invitrogen, 11668019) following the manufacturer’s instructions. Briefly, 1.5 μg of total DNA (for protein expression analyses) or 2 μg GluN1, 2 μg HA-MDGAs, and 4 μg GluN2A or 2B (for Co-IPs), were mixed with Lipofectamine at a 1:1 ratio in 100 μL of pre-warmed Opti-MEM (Gibco, 31985062) for 15 minutes (min) at room temperature (RT). The resulting mixture was added to an individual well of a 6-well plate. When GluN2A or 2B was co-transfected with GluN1, 50 μM AP-5 and 20 mM MgCl2 were added 4 hours (hr) after transfection. 24-48 h post transfection, cells were washed and collected in ice-cold PBS and centrifuged.

### Co-immunoprecipitation and immunoblotting

For Western blot, after centrifugation, pelleted cells were lysed with 200 μL of 4x SDS-PAGE sample buffer, sonicated, denatured for 5 min at 65°C, and centrifuged at 20,000 x g for 5 min to pellet insoluble cellular debris. Protein lysates (5 μL or 2.5% of sample, avoiding the pelleted debris) were separated by SDS-PAGE. For more details see (*47*).

For co-IP, after centrifugation, cell pellets were resuspended in 1% Triton X-100 lysis buffer and lysed for 1 hr at 4°C. Protein lysates were collected after centrifugation. Then, the HA antibody (Cell Signaling #3724) was added to lysates and incubated overnight at 4°C. Next day, protein A Sepharose (Sigma-Aldrich #P3391) was added to the mixture and incubated for 4 hr at 4°C. The Protein A Sepharose-attached antibody-protein complexes were washed with 1% Triton X-100 lysis buffer 3 times. 2X SDS sample buffer was added to the complexes and incubated at 42°C for 20 min for protein elution. The eluted proteins were separated with 7% SDS-PAGE.

WT and HA-MDGA1/Myc-MDGA2 brain tissue was processed as previously described (*48*). Briefly, either the entire forebrain or dissected brain regions, were collected in ice-cold PBS and homogenized in buffer containing 20 mM Tris-HCl (pH 7.5), 0.32 M sucrose (Millipore Sigma, 573113), 5 mM EDTA (Sigma-Aldrich, 6381-92-6) and protease and phosphatase inhibitors (Roche, #11836170001). After centrifugation at 1,000g for 10 min to remove the nuclear fraction, the supernatant (S1) was either mixed with SDS-containing sample buffer or centrifuged at 10,000g for 15 min to obtain the P2 fraction. The P2 fraction was then re- suspended in SDS-containing sample buffer.

All samples were assessed by PAGE-SDS electrophoresis. Immuno-Blot® PVDF membranes (Bio-Rad, #1620177) were blocked with 5% blotting grade nonfat milk (Lab Scientific, #M0841) in tris buffered saline buffer with 0.1% tween 20 (Sigma-Aldrich, #P1379). The following primary antibodies (company, cat no.) were used in Western blot experiments: mouse anti-beta-actin (ABM, #G043), mouse anti-Flag (Sigma, #F1804), mouse anti-GluN1 (Thermo #32-0500), mouse anti-HA (Roche, #11 867 423 001), rabbit anti-HA (Cell Signaling, #3724), rabbit anti-Myc (Abcam, #ab9106), mouse anti-α-tubulin (Sigma-Aldrich, #T9026).

HRP-conjugated secondary antibodies raised against the appropriate species were used: anti-mouse IgG (GE Healthcare, #NA931), anti-rabbit IgG (GE Healthcare, #NA934), anti-rat IgG (Cell Signaling Technology, #7077), Anti-Rabbit IgG (Vector laboratories #PI-1000), anti-mouse IgG (Vector laboratories #PI-2000). Clarity^TM^ Western ECL (BioRad, #170-5060) was then added to membranes. Western blots were imaged using BioRad Chemidoc. Blot images were analysed by creating a uniformly sized box around each desired band, leaving room above and below. A histogram measuring the band intensity across the length of the box was created. A level base for the histogram was then approximated, encompassing an area containing the lowest signal point to the highest signal point. The area of the resulting triangle was then measured via the FIJI (Image J, Janelia) software. To represent the developmental time-course of MDGA expression, HA-MDGA1 / α-tubulin and Myc-MDGA2 / α-tubulin ratios were calculated and normalized to a reference age (P3). All data are presented as mean ± SEM.

### Immunofluorescence

For immunofluorescence analyses, 4% paraformaldehyde (PFA) fixed coronal or sagittal brain slices (30 μm thick) were processed. After blocking tissue with 5% swine serum and 2% BSA in permeabilizing conditions (0.1% Triton X-100, Sigma-Aldrich, # T8787), immunofluorescence was performed by overnight incubation at 4 °C with a rabbit anti-HA primary antibody (Cell Signaling, #3724, 1:500 dilution), in combination with one of four primary guinea pig antibodies against synaptic markers: Homer1b/c (Synaptic Systems, #160 025, 1:200), Nlgn2 (Synaptic Systems, #129 205, 1:500), vGluT1 (Synaptic Systems, #135 304, 1:15,000), vGAT (Synaptic Systems, #129 205, 1:500). This was followed by incubation with corresponding Alexa 488 goat anti-guinea pig (Invitrogen, #A-11073, 1:500) and Alexa 594 goat anti-rabbit (Invitrogen, #A-11012, 1:500) secondary antibodies. Slides were mounted with either Vectashield Antifade Mounting Medium with DAPI (Vector Laboratories, # H-1200) or ProLong Gold Antifade Reagent with DAPI (Cell Signaling Technology, # 8961S).

### Fluorescence Tomography microscopy and quantification

Image z-stacks of hippocampal field CA1 stratum radiatum (SR) were collected at 0.2 µm steps using a 1.4 NA 63X objective on a Leica DM6000 epifluorescence microscope equipped with a Hamamatsu ORCA-ER digital camera. The image sample field size was 105 x 136 (x,y) with a 2 µm depth (z), for a total size of 28,560 µm^3^. For each antisera combination, 6- 8 image stacks from three sections per brain were taken. On average, the total numbers of synaptic profiles assessed per image stack in the P15 brain were: 9,715 ± 1,236 (s.e.) for Homer, 6,515 ± 325 for Nlgn2, 16,950 ± 1,651 for vGluT1, and 10,674 ± 280 for vGAT. The greater number of excitatory versus inhibitory synaptic profiles at this age is consistent with previous work showing greater asymmetric versus symmetric shaft synapses on CA1 stratum pyramidale (SP) neurons (Watson et al. 2016).

For quantification of immunofluorescent-labeled puncta, image stacks were pre- processed by standardizing the dynamic range via inserting two high and two low intensity reference squares (100x100 pixels; 1µm^2^) to the green and red channels of all images (Python 3.8 with NumPy, skimage.io, os, PIL, tifffile, and json). This step allows the subsequent quantification of all puncta intensity to be normalized to the global reference square intensity for each channel, rather than the maximum intensity within the image, without largely altering the background or existing raw pixel values. Image stacks were then analyzed using in-house software to quantify double-labeled, single-labeled, and all-labeled puncta within the size constraints of synapses as previously described (*49–52*). Background staining variations in the deconvolved images were normalized to 30% of maximum background intensity using a Gaussian filter. Object recognition and measurements of immunolabeled puncta were automated using software built in-house using Matlab R2019b, Perl, and C which allows for detailed analysis of objects reconstructed in 3 dimensions (3D). Pixel values (8-bit) for each image were multiply binarized using a fixed-interval intensity threshold series followed by erosion and dilation filtering to reliably detect edges of both faintly and densely labeled structures. Object area and eccentricity criteria were applied to eliminate from quantification elements that do not fit the size and shape range of synaptic structures, including the global reference squares. For synaptic compartment localization of the HA-tag, immunolabeled puncta were considered colocalized if they touched or overlapped to any degree as assessed in 3D. Immunolabeled object counts were averaged across sections to produce mean values for each measure per animal.

### Confocal Imaging and Image Analysis

Brain sections were imaged with a Leica Sp8 confocal microscope (Leica Microsystems, Wetzlar, Germany) equipped with six single photo laser lines (405 nm, 458, 488, 514, 568, and 633 nm) and four detectors at the University of California, Irvine Optical Biology Core. Images were acquired using a 63x oil objective as a series of 20 z-steps, with a z-step size of 1.38 μm, at a resolution of 1024 x 1024 pixels, scanning frequency of 400 Hz. The optical resolution (voxel size) per image was 180 nm in the xy-plane and 1.38 μm in the z-plane. Images were saved in a “lif.” format, and analysis and quantification of total synaptic puncta and puncta colocalization was performed using Imaris 9.9.1 (Bitplane, South Windsor, CT, USA) and MatLab Runtime R2022b (Mathworks, Natick, MA, USA).

Imaris analysis entailed a software-specific conversion of the original “lif.” file into an “ims.” format, allowing for three-dimensional analysis. The new “ims.” file contained both the original image and its stored metadata. Generally, the “Spots” tool was utilized to assign representative three-dimensional ellipsoid shapes to cover individual puncta. This included puncta of each of the four synaptic markers (Homer1b/c, Nlgn2, vGluT1, and vGAT) as well as HA-MDGA1. These spots were then used as a proxy for synaptic puncta during further colocalization analysis and quantification. Once in Imaris, the brightness and contrast settings for CH2 (HA-MDGA1, Alexa 594) and CH3 (Synaptic marker, Alexa 488) were adjusted to qualitatively minimize background noise in each channel and emphasize specific signal. These settings were then applied across all KI and WT images. When creating the “spots” for HA- MDGA1 and each of the synaptic markers, the following protocol was followed. First, the minimum xy and z diameters of HA-MDGA1 puncta were set to 0.5 μm and 0.9 μm, respectively. The same was done with dimensions of 0.4 µm in the xy-plane and 0.9 in the z-plane for the synaptic markers Homer1b/c, Nlgn2, and vGAT, and 0.5 µm and 0.9 µm in the xy- and z-planes, respectively, for vGluT1. The “Background Subtraction” option available when creating spots was then used. Technically, this option smooths the image prior to the addition of spots by using a Gaussian filtered channel (Gaussian filtered by ¾) minus the intensity of the original channel Gaussian filtered by 8/9th of the punctum radius. A region of interest (ROI) was created to restrict the colocalization quantification to solely within SR of each image. This ROI spanned an average area of approximately 15,9812±1506 μm^2^ across all samples. When building the corresponding representative spots, the number of spots were adjusted qualitatively using the automatically generated and interactive “Quality” filter histogram to select what appeared to be specific dense signal while excluding faint puncta that appeared to be background signal. To ensure an accurate spot segmentation of the underlying puncta determined by size, the “Different Spots Sizes” selection was utilized. Within this setting, the “Local Contrast” tool was used. The corresponding histogram was manually adjusted to ensure each spot covered as much of the puncta as possible. Spots were then rendered. Once optimal settings for each of these parameters were established for HA-MDGA1 and each of the four synaptic markers, a batched protocol to automate spots creation on every image was run. Despite efforts to minimize non-specific HA signal, some residual and dim HA puncta were still detected in WT samples (Fig. 1J). Therefore, the Imaris filter selection tool: “Intensity Max”, was applied, setting the reference value to the 10% highest intensity spots on a HA-MDGA1 KI sample, and discarding spots with intensity values below the threshold, thereby allowing for a standardized level of comparison between samples. As expected, this processing resulted in comparatively less colocalizing spots in WT samples compared to the KI samples (Fig. 2). To determine the colocalization between HA- MDGA1 and synaptic marker spots, the Matlab extension “Spots Colocalize” was used. This extension determines colocalization between two or more spots found within a given length measured from the center of each spot. Because both HA-MDGA1 and synaptic marker puncta appeared at varying sizes, setting the colocalization parameter to consider only spots at or within 0.3 μm from the center of neighbouring puncta was found to be the most accurate. Spots colocalizations were reported as the number of discrete HA-MDGA1 spots colocalized with at least one spot of each corresponding synaptic marker per 100 square microns.

### Attempts to minimize artefacts in tagged protein quantification

We found a variable amount of residual, punctate HA expression in CA1 in WT mice, which represented approximately 30% of the signal found in the HA-MDGA1 mice. For this reason, KI mice and WT counterparts were compared in all imaging experiments. In any case, our experience constitutes a cautionary note when characterizing the expression of tagged proteins using KI mice.

### Spine morphology measurements

Images were acquired using super-resolution microscopy (N-SIM Microscope System, Nikon) in organotypic slice preparations at day in vitro (DIV) 7, after sparse transfection of MDGA1 and MDGA2 shRNAs or control shRNA together with GFP at DIV 1. For use with the available inverted microscope and oil-immersion objective lens, slices were fixed in 4% PFA/4% sucrose in PBS and washed 3 × with PBS. To amplify the GFP signal, slices were then blocked and permeabilized in 3% BSA in PBS containing 0.1% Triton X-100 (Sigma-Aldrich # T8787) and stained with rabbit anti-GFP (2 μg/mL, Life Technologies, #A-11122) followed by washes in PBS-Tx and staining with Alexa 488-conjugated goat anti-rabbit (4 μg/mL, Life Technologies, #A11034). Slices were then mounted in SlowFade Gold (Life Technologies, #S36936) for imaging. Only dendrites in the top 20 μm of the slice were imaged. Some slices were further processed with an abbreviated SeeDB-based protocol (*53*) in an attempt to reduce spherical aberration, but no substantial improvement was seen. Images were acquired with a ×100 oil objective in 3D-SIM mode using supplied SIM grating (3D EX V-R ×100/1.49) and processed and reconstructed using supplied software (NIS Elements, Nikon). Morphological analysis was done on individual sections using ImageJ to perform geometric measurements on spines extending laterally from the dendrite. Spine neck widths were obtained from full width half-maximum measurements based on Gaussian fits of line profile plots (*54*). Neck length was measured from the base of the spine to the base of the head. Head diameter was measured perpendicular to the spine neck axis through the thickest part of the spine head, and diameter was obtained using full width tenth maximum (FWTM) measurements based on Gaussian fits to approximate manual head measurement.

### Protein Identification Using Reversed-phase Liquid Chromatography Electrospray Tandem Mass Spectrometry (LC-MS/MS)

HA-tagged MDGA1 or MDGA2 were expressed in HEK293T (as described above) and two days post-transfection were released from the membrane by addition of phospholipase C (PLC). Briefly, for PLC treatment, transfected 293T cell monolayers were washed, and then incubated with 0.2U/ml PLC (Sigma-Aldrich, P7633) in Optimem for 2 hr at 37°C. Soluble MDGAs were purified from the cellular media with anti-HA antibodies (see above) and incubated with mouse brain P2 fractions (solubilized in 1% triton) overnight to identify interacting proteins. Pull-downs were separated on SDS-page gels and subjected to silver staining (Thermo Scientific, *#*24612). See Fig. 3A for more details. Post silver staining, the targeted gel bands were excised from a gel and subjected to in-gel tryptic digestion. Proteins in the gel band were reduced with 10 mM dithiothreitol in 25 mM ammonium bicarbonate at 56°C for 1 hr, followed by alkylation with 55 mM iodoacetamide in 25 mM ammonium bicarbonate at room temperature for 45 min in the dark. The samples were then incubated overnight at 37°C with 100 ng trypsin (sequence grade, Promega). The peptides formed from the digestion were further purified by µC18 ZipTips (Millipore) and resuspended in 0.1% formic acid in HPLC water.

The LC–MS/MS analyses were conducted by either a Velos Pro Elite Orbitrap (Elite) Mass Spectrometer or a LTQ Orbitrap Velos (Velos) mass spectrometer (Thermo Scientific) coupled with a NanoAcquity UPLC system (Waters). During the LC separation, peptides were first loaded onto an Easy-Spray PepMap column (75 μm x 15 cm, Thermo Scientific). Following the initial column equilibration in 98% A (0.1% formic acid in water) / 2% B (0.1% formic acid in acetonitrile) over 20 min, the concentration of the phase B was linearly increased from 2 – 30% at a flow rate of 300 nL per min over 27 min. Then the phase B concentration was increased linearly from 30 – 50% sequentially in the next two min. The column was then re-equilibrated in 98% A / 2% B over 11 min. After a survey scan in the Orbitrap, the top six most intensive precursor ions were fragmented by either collision-induced dissociation with the Elite or Higher-energy C-trap dissociation with the Velos. The acquired MS/MS raw data was converted into peaklists using an in-house software PAVA (*55*). The peaklists were then searched against the Uniprot Mus Musculus database (UniProtKB.2017.11.01) using Protein Prospector search engine (http://prospector.ucsf.edu/prospector/mshome.htm). Proteins with at least 1 unique peptide were reported.

Mass spectrometry experiments were repeated three independent times for each condition (control (GFP), MDGA1, and MDGA2). Proteins were considered binders if i) they were identified >1 experiment and ii) 0 peptides were identified in the control lanes. Binders are highlighted in yellow in Supplementary Table 1. Fig. 3B contains a shortened list of synaptic proteins of interest.

### Electrophysiology

Hippocampal organotypic slice cultures were isolated from P6-8 rats, as described previously (*56*) and biolistically transfected at DIV1. Briefly, mixed plasmid DNA (50 μg total) was coated on 1 μM-diameter gold particles (Bio-Rad, 1652263) in 0.5 mM spermidine. The DNA was precipitated with 0.1 mM CaCl2, washed four times in ethanol (Sigma-Aldrich, 459836) and coated onto PVC tubing (Bio-Rad, 1652441). The tubing was dried with N2 gas, and the DNA-coated gold particles were delivered to the slices with a Helios Gene Gun (BioRad). Equal amounts of plasmid DNA (when necessary) were coated to gold particles for excitatory and inhibitory recordings, respectively. Each plasmid expressed different fluorescent markers, and we only recorded from cells that expressed both GFP and mCherry signifying expression of both plasmids. Slices were maintained at 34°C with media changes every other day.

Dual whole-cell recordings from CA1 PNs were performed at DIV 7-9. Since biolistics results in sparsely transfected hippocampal PNs per slice, simultaneous recordings from both a transfected neuron and neighboring untransfected control neuron were collected. Synaptic responses were evoked by stimulating with a monopolar glass electrode filled with artificial cerebrospinal fluid (aCSF) in CA1 SR. Synaptic strength was calculated by comparing the difference in magnitude of the transfected cell to the non-transfected control cell recorded simultaneously. PNs were identified by morphology and location. The number of experiments (n) reported in the figure legends refer to the number of paired recordings. Membrane holding current, input resistance, and pipette series resistance were monitored throughout recording. All recordings were made at 20-25°C using glass patch electrodes filled with an internal solution consisting of 135 mM CsMeSO4 (Sigma-Aldrich, C1426), 8 mM NaCl (Sigma-Aldrich, 7647-14-5), 10 mM HEPES, 0.3 mM EGTA (Sigma-Aldrich, E3889), 4 mM Mg-ATP (Sigma-Aldrich, A9187), 0.3 mM Na-GTP (Sigma-Aldrich, G8877), 5 mM QX-314 (Abcam, 5369-03-9), and 0.1 mM spermine (Sigma-Aldrich, S2876), and an external solution containing 119 mM NaCl, 2.5 mM KCl (Sigma-Aldrich, 60128), 4 mM MgSO4 (Sigma-Aldrich, 63138), 4 mM CaCl2, 1 mM NaH2PO4 (Sigma-Aldrich, S9638), 26.2 mM NaHCO3 (Sigma-Aldrich, S8875) and 11 mM glucose (Sigma-Aldrich, G8270) bubbled continuously with 95% O2 and 5% CO2. Excitatory recordings were made in the presence of 100 μM picrotoxin (TCI, C0375) to block inhibitory currents and a small (50 nM) amount of NBQX (abcam, ab120046) to reduce epileptiform activity at -70 mV (AMPA). Inhibitory recordings were made in the presence of 100 μM D-APV (Alomone Labs, D-145) and 10 μM NBQX to block NMDA receptor (NMDAR)- and AMPAR-mediated currents, respectively, at 0 mV. AMPAR- and IPSC-mediated currents were measured at the peak of the current. Investigator was blinded to the control vs. experimental group during data analysis. For more details see (*47*).

### Statistics

Graph Pad Prism 9 was used for analyses of statistical significance and outliers.

Statistical significance of immunoblots was tested using a Mann-Whitney U test. Paired whole-cell recordings were analyzed with a Wilcoxon signed-rank test. Unpaired Student’s T-Test, and one- or two-way ANOVA followed by Tukey’s post hoc test for multiple comparisons were used as appropriate to compare experimental groups in synaptic colocalization analyses. For the confocal HA-vGluT1 colocalization analysis (Fig. 2G, H), three HA-MDGA1 brains were excluded as image acquisition settings for those samples were found to not match those of the rest of the samples analysed. All data are presented as mean ± SEM.

## Supporting information

Combined supplementary figures

## Acknowledgements

We would like to thank Eric Dang, Dan Qin and Ananth V. Kolli for excellent technical assistance and members of the Nicoll and Diaz Alonso lab for helpful comments throughout the project. Funding sources: K99/R00 MH118425, BBRF Young Investigator Award 30264 and UCI Institutional funding to J.D.-A., R01 MH117139 and R01 MH070957 to R.A.N. and R01 HD101642 to C.M.G. Mass Spectrometry of this work was provided by the Mass Spectrometry Resource at UCSF (A.L. Burlingame, Director) supported by the Dr. Miriam and Sheldon G. Adelson Medical Research Foundation (AMRF).

## Author contributions

M.A.B., M.S., A.A.L., S.W., V.N.C., J. L., S.I., K.H.L. and J.D.-A., performed experiments; M.A.B., M.S., A.A.L., S.W., V.N.C., J. L., S.I., K.H.L., A.L.B., K.R., C.M.G., R.A.N and J.D.- A. analyzed data; M.A.B., J.D.-A. and R.A.N designed research; M.A.B., M.S., J. L., C.M.G., R.A.N and J.D-A drafted and all authors commented on the manuscript and R.A.N. and J.D.-A. coordinated research.

## Data and materials availability

All data needed to reproduce and evaluate the conclusions of the study are present in the paper, including the supplementary materials. Materials will be available upon completion of the appropriate MTAs.

