## Supplementary material for "Contrasting synaptic roles of MDGA1 and MDGA2": Combined supplementary figures

### Figure S1

#### A CRISPR/Cas9 homology directed repair based HA-MDGA1 KI strategy overview

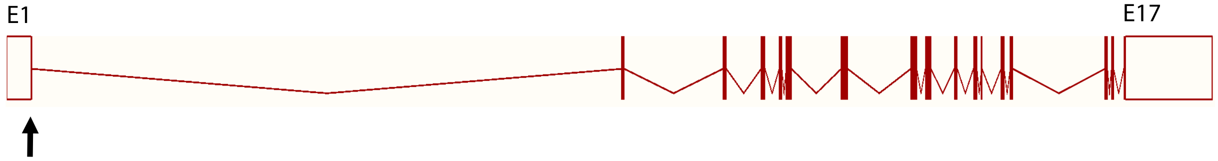


Cas9 cut site

WT *MDGA1* sequence 5' TATCCCCTTCCACTGTCGGGGACAAGGAGTCT 3'

gRNA sequence (targeting forward strand) 5' CCCUUCCACUGUCGGGGACAAGG 3'

Inserted HA-linker sequence 5' TACCCATACGATGTTCCAGATTACGCTAGATCTCGAGAT 3'

*HA-MDGA1* sequence 5' TATCCCCTTCCACTGTCGGGGATACCCATACGATGTTCCAGATTACGCTAGATCTCGAGATCAAGGAGTCT 3'

#### B HA-MDGA1 heterozygous founder mouse sequence

KI strand sequence TACCCATTACGATGTTCCAGATTACGCTAGATCTCGAGATCAAGGAGTCTATGGTAAGAGCCAG

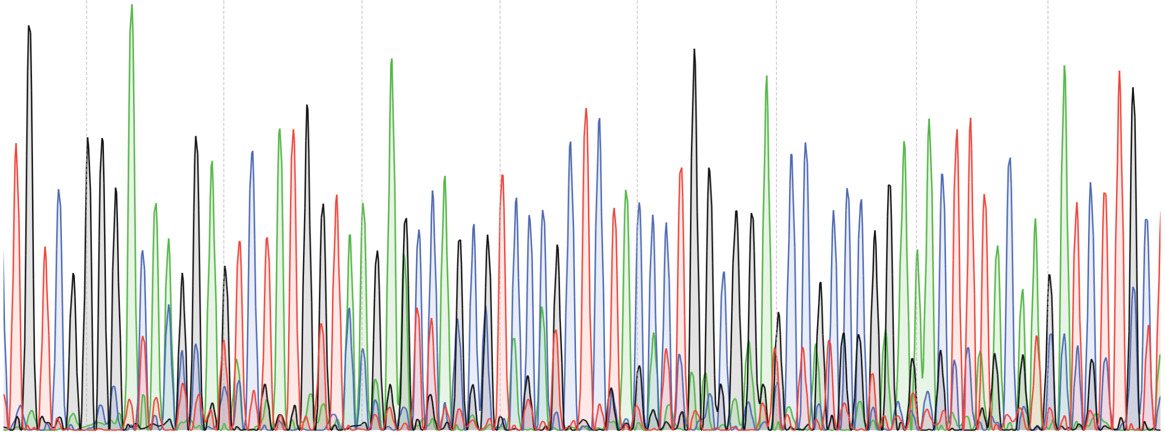

#### C Predicted protein sequence

1 MEVTCLLLLA LIPFHCRG**YP YDVPDYARSR D**QGVYAPAQA QIVHAGQACV
51 VKEDNISERV YTIRESDTLV LQCLVTGHPR PQVRWTKTAG SASDKFQETS
101 VFNETLRIER IARTQGGRYY CKAENGVGVP AIKSIRVDVQ YLDEPVLTVH
151 QTVSDVRGNF YQEKTVFLRC TVSSNPPARF IWKRGSDTLS HSQDNGVDIY
201 EPLYTQGETK VLKLKNLRPQ DYASYTCQVS VRNVCGIPDK AITFQLTNTT
251 APPALKLSVN ETLVVNPGEN VTVQCLLTGG DPLPQLHWSH GPGPLPLGAL
301 AQGGTLSIPS VQARDSGYYN CTATNNVGNP AKKTVNLLVR SLKNATFQIT
351 PDMIKESENI QLGQDLKLSC HVDAVPQEKV NYQWFKNGKP ARTSKRLLVT
401 RNDPELPAVT SSLELIDLHF SDYGTYLCMA SFPGSPVPDL SIEVNISSET
451 VPPTISVPKG RAVVTVREGS PAELQCEVRG KPRPPVLWSR VDKEAALLPS
501 GLALEETPDG KLRLESVSRD MSGTYRCQTA RYNGFNVRPR EAQVQLTVHF
551 PPEVEPSSQD VRQALGRPVL LRCSLLRGSP QRIASAVWRF KGQLLPPPPV
601 LPAAAVETPD HAELRLDALT RDSSGNYECS VSNDVGSATC LFQVSAKAYS
651 PEFYFDTPNP TRSHKLSKNY SYVLQWTQRE PDAVDPVLNY RLSIRQLNQH
701 NAMVKAIPVR RVEKGQLLEY ILTDLRVPHS YEIRLTPYTT FGAGDMASRI
751 IHYTEHNTCH FEDEKICGYT QDLTDNFDWT RQNALTQNPK RSPNTGPPTD
801 ISGTPEGYYM FIETSRPREL GDRARLVSPL YNASAKFYCV SFFYHMYGKH
851 IGSLNLLVRS RNKGTLDTHA WSLSGNKGNV WQQAHVPINP SGPFQIIFEG
901 VRGSGYLGDI AIDDVTLKKG ECPRRQMDPN KVVVMPGSGA PRLSSLQLWG
951 SMAIFLLALQ R

### Figure S2

**A** CRISPR/Cas9 homology directed repair based Myc-MDGA2 KI strategy overview

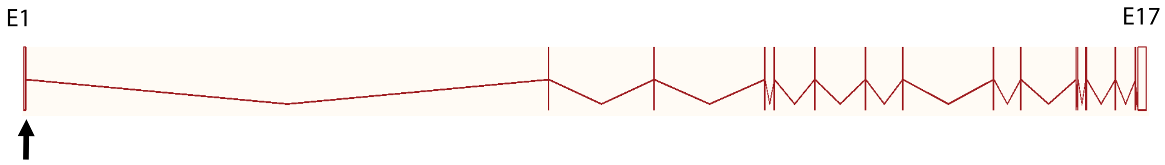


Cas9 cut site

intron

WT *MDGA2* sequence 5' GAGGGAATCTCTGGCCAAGGAGTGTACGGTGAGTGGAATC 3'

gRNA sequence (targeting reverse strand) 5' TCCACTCACCGTACACTCCTTGG 3'

Inserted Myc-linker sequence 5' GAACAAAAACTCATCTCAGAAGAGGATCTGAGATCTCGAGAT 3'

Myc-MDGA2 sequence 5' GAGGGAATCTCTGGCGAACAAAAACTCATCTCAGAAGAGGATCTGAGATCTCGAGATCAAGGAGTGTACGGTGA 3'

**B** Myc-MDGA2 heterozygous founder mouse sequence (reverse strand)

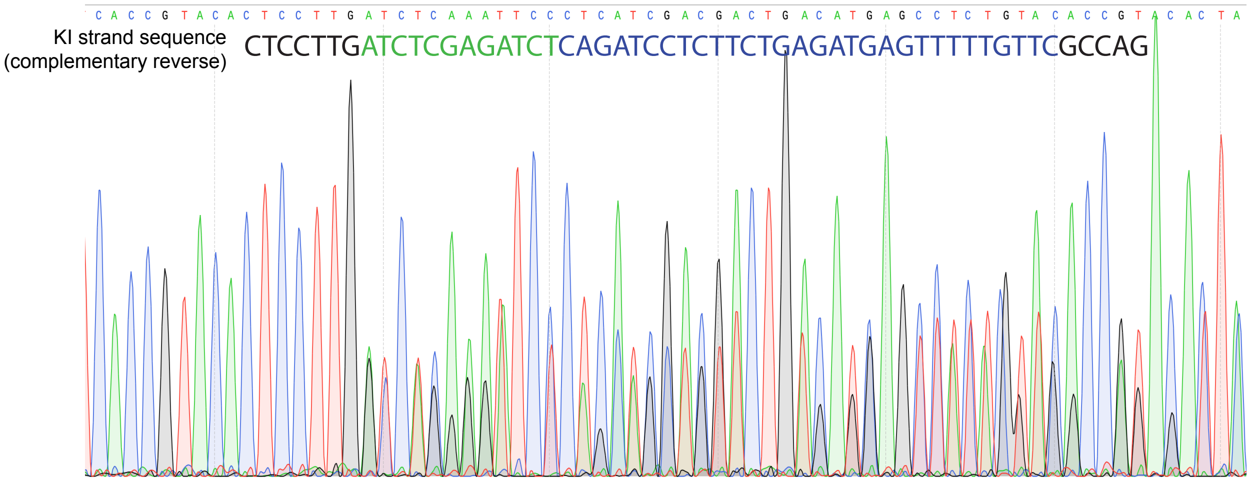


**C** Predicted protein sequence

1 MDLVYGLVWL LTVLLEGISG EQKLISEEDL RSRDQGVYAP PTVRIVHSGL
51 ACNIEEERYS ERVYTIREGE TLELTCLVTG HPRPQIRWTK TAGSASDRFQ
101 DSSVFNETLR ITNIQRHQGG RYYCKAENGL GSPAIKSIRV DVYYLDDPVV
151 TVHQSIGEAK EQFYYERTVF LRCVANSNPP VRYSWRRGQE VLLQGSDKGV
201 EIYEPFFTQG ETKILKLKNL RPQDYANYSC IASVRNVCNI PDKMVSFRLS
251 NKTASPSIKL LVDDPIVVNP GEAITLVCVT TGGEPTPSLT WVRSFGTLPE
301 KIVLNGGTLT IPAITSDDAG TYSCIANNNV GNPAKKSTNI IVRALKKGRF
351 WITPDPYHKD DNIQIGREVK ISCQVEAVPS EELTFSWFKN GRPLRSSERM
401 VITQTDPDVS PGTTNLDIID LKFTDFGTYT CVASLKGGGI SDISIDVNIS
451 SSTVPPNLTV PQEKSPLVTR EGDTIELQCQ VTGKPKPIIL WSRADKEVAM
501 PDGTMQMESY DGTLRIVNVS REMSGMYRCQ TSQYNGFNVK PREALVQLIV
551 QYPPAVEPAF LEIRQGQDRS VTMSCRVLRA YPIRVLTYEW RLGNKLLRTG
601 QFDSQEYTEY PLKSLSNENY GVYNCSIINE AGAGRCSFLV TGKAYAPEFY
651 YDTYNPVWQN RHRVYSYSLQ WTQMNPDAVD RIVAYRLGIR QAGQQRWWEQ
701 EIKINGNIQK GELITYNLTE LIKPEAYEVR LTPLTKFGEG DSTIRVIKYT
751 APVNPHLREF HCGFEDGNIC LFTQDDTDNF DWTKQSTATR NTKYTPNTGP
801 SADRSGSKEG FYMYIETSRP RLEGEKARLL SPVFSIAPKN PYGPTNSAYC
851 FSFFYHMYGQ HIGVLNVYLR LKGQTTIENP LWSSSGNKGQ RWNEAHVNIY
901 PITSFQLIFE GIRGPGIEGD IAIDDVSIAE GECAKQDLPT KNSVDGAVGI
951 LVHIWLFPVI ILISILSPRR

Figure S3

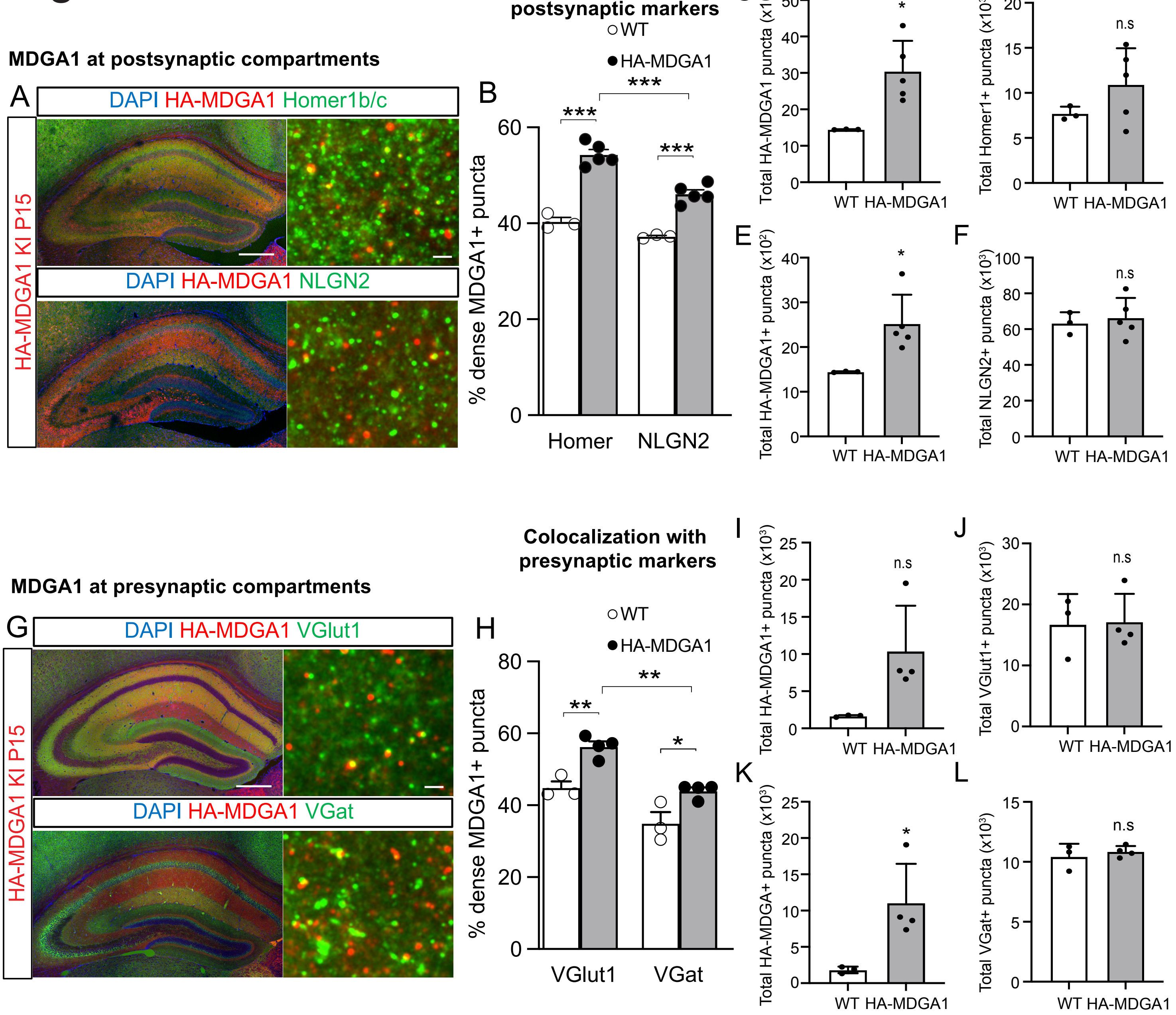
MDGA1 at postsynaptic compartments
A
DAPI HA-MDGA1 Homer1b/c
DAPI HA-MDGA1 NLGN2
HA-MDGA1 KI P15
Colocalization with postsynaptic markers
WT
HA-MDGA1
B
% dense MDGA1+ puncta
Homer
NLGN2
C
Total HA-MDGA1 puncta (x10²)
D
Total Homer1+ puncta (x10³)
E
Total HA-MDGA1+ puncta (x10²)
F
Total NLGN2+ puncta (x10³)
WT HA-MDGA1
n.s
MDGA1 at presynaptic compartments
G
DAPI HA-MDGA1 VGlut1
DAPI HA-MDGA1 VGat
HA-MDGA1 KI P15
Colocalization with presynaptic markers
H
% dense MDGA1+ puncta
VGlut1
VGat
I
Total HA-MDGA1+ puncta (x10³)
J
Total VGlut1+ puncta (x10³)
K
Total HA-MDGA+ puncta (x10³)
L
Total VGat+ puncta (x10³)

### Figure S4

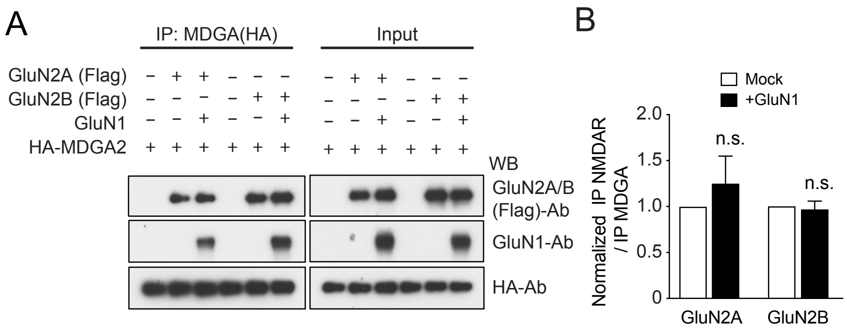
A
IP: MDGA(HA)
Input
GluN2A (Flag)
GluN2B (Flag)
GluN1
HA-MDGA2
WB
GluN2A/B (Flag)-Ab
GluN1-Ab
HA-Ab
B
Mock
+GluN1
Normalized IP NMDAR / IP MDGA
2.0
1.5
1.0
0.5
0
n.s.
n.s.
GluN2A
GluN2B

### Figure S5

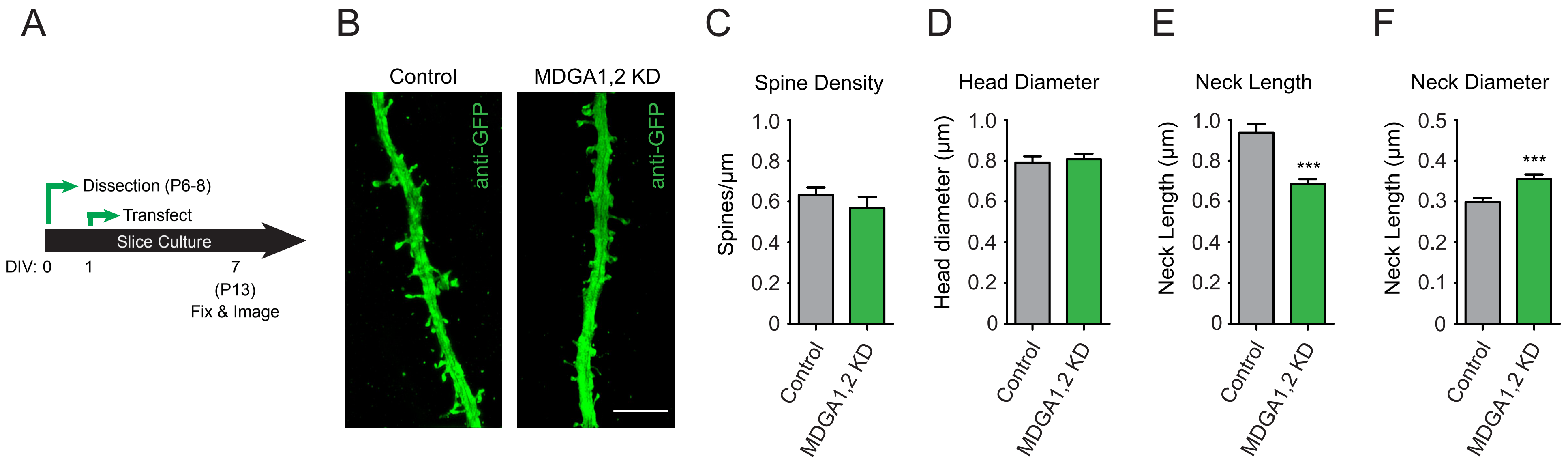

A
Dissection (P6-8)
Transfect
Slice Culture
DIV: 0
1
7
(P13)
Fix & Image
B
Control
MDGA1,2 KD
anti-GFP
anti-GFP
C
Spine Density
Spines/µm
1.0
0.8
0.6
0.4
0.2
0
Control
MDGA1,2 KD
D
Head Diameter
Head diameter (µm)
1.0
0.8
0.6
0.4
0.2
0
Control
MDGA1,2 KD
E
Neck Length
Neck Length (µm)
1.0
0.8
0.6
0.4
0.2
0
***
Control
MDGA1,2 KD
F
Neck Diameter
Neck Length (µm)
0.5
0.4
0.3
0.2
0.1
0
***
Control
MDGA1,2 KD

### Figure S6

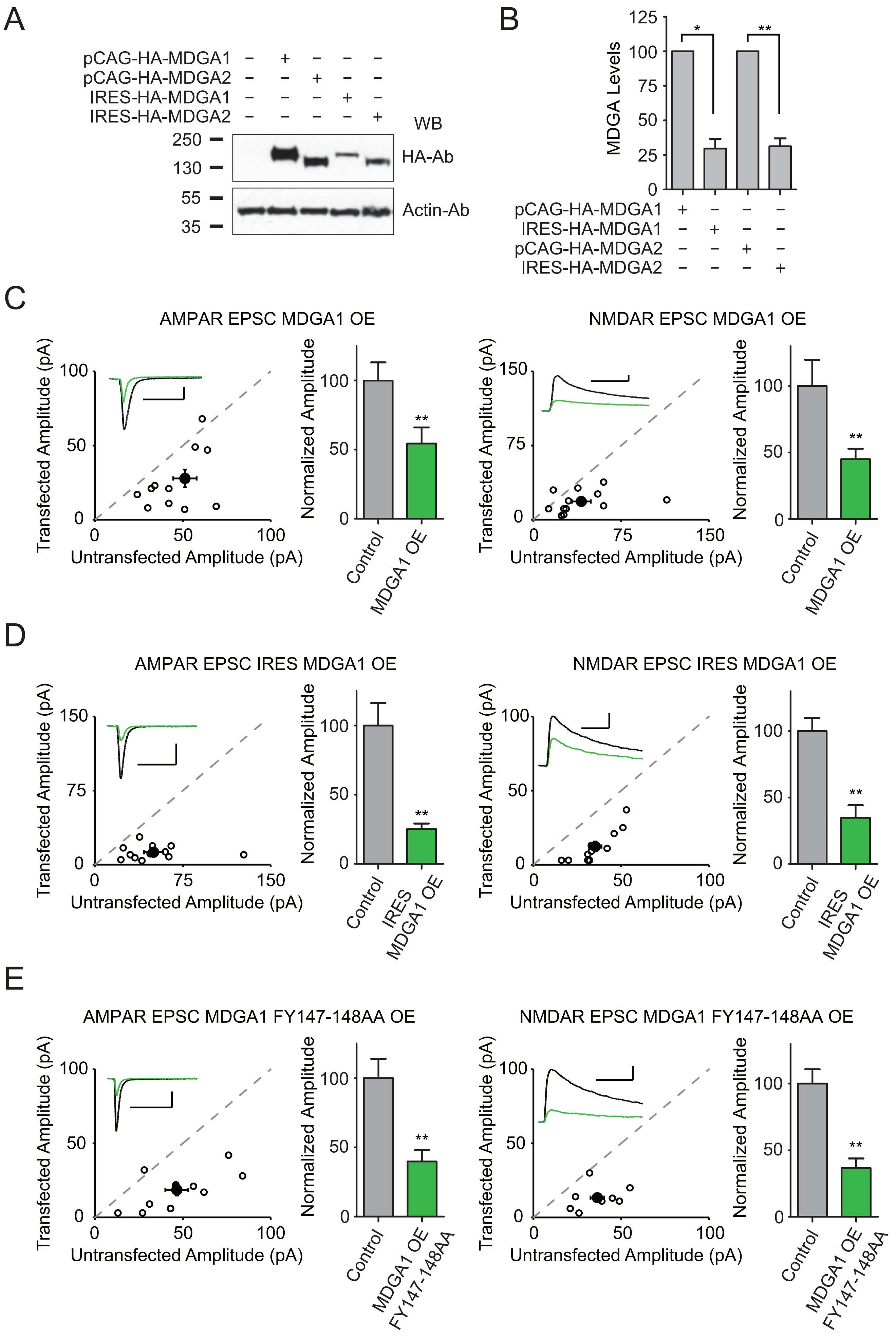

A
pCAG-HA-MDGA1 – + – – –
pCAG-HA-MDGA2 – – + – –
IRES-HA-MDGA1 – – – + –
IRES-HA-MDGA2 – – – – +
WB
250
130
HA-Ab
55
35
Actin-Ab
B
MDGA Levels
125
100
75
50
25
0
*
**
pCAG-HA-MDGA1 + – – –
IRES-HA-MDGA1 – + – –
pCAG-HA-MDGA2 – – + –
IRES-HA-MDGA2 – – – +
C
AMPAR EPSC MDGA1 OE
NMDAR EPSC MDGA1 OE
Transfected Amplitude (pA)
Untransfected Amplitude (pA)
Normalized Amplitude
Control
MDGA1 OE
**
D
AMPAR EPSC IRES MDGA1 OE
NMDAR EPSC IRES MDGA1 OE
Transfected Amplitude (pA)
Untransfected Amplitude (pA)
Normalized Amplitude
Control
IRES MDGA1 OE
**
E
AMPAR EPSC MDGA1 FY147-148AA OE
NMDAR EPSC MDGA1 FY147-148AA OE
Transfected Amplitude (pA)
Untransfected Amplitude (pA)
Normalized Amplitude
Control
MDGA1 OE FY147-148AA
**

### Figure S7

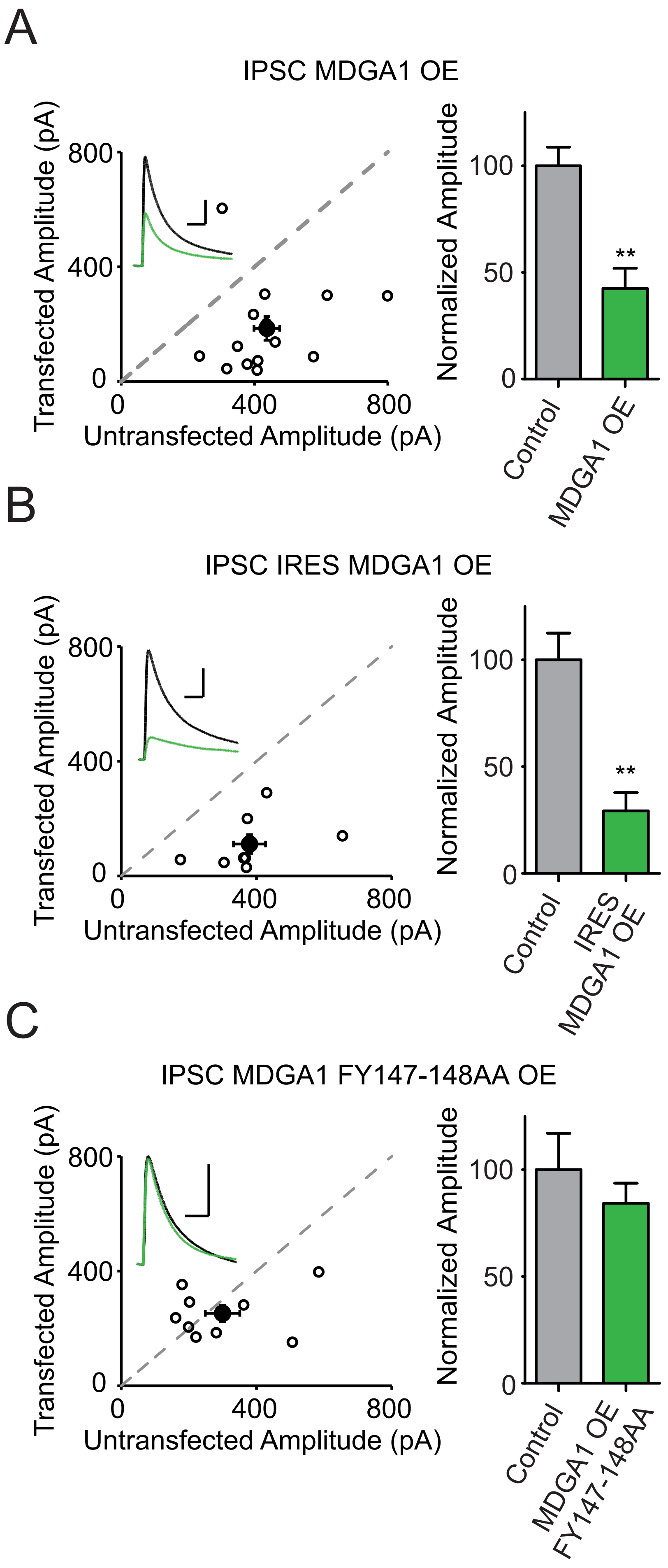
A
IPSC MDGA1 OE
Transfected Amplitude (pA)
800
400
0
0
400
800
Untransfected Amplitude (pA)
Normalized Amplitude
100
50
0
**
Control
MDGA1 OE
B
IPSC IRES MDGA1 OE
Transfected Amplitude (pA)
800
400
0
0
400
800
Untransfected Amplitude (pA)
Normalized Amplitude
100
50
0
**
Control
IRES MDGA1 OE
C
IPSC MDGA1 FY147-148AA OE
Transfected Amplitude (pA)
800
400
0
0
400
800
Untransfected Amplitude (pA)
Normalized Amplitude
100
50
0
Control
MDGA1 OE FY147-148AA

**Fig S1. Generation of HA-MDGA1 mice.** A, Strategy for CRISPR/Cas9 homology-directed repair mediated HA tag insertion in the MDGA1 locus (image adapted from *Ensembl genome browser*). B, Representative sequencing of the MDGA1 locus after successful insertion of the HA tag in the HA-MDGA1 KI colony founder mouse. C, Predicted translation of the MDGA1 locus after successful insertion of the HA tag.

**Fig S2. Generation of Myc-MDGA2 mice.** A, Strategy for CRISPR/Cas9 homology-directed repair mediated Myc tag insertion in the MDGA2 locus (image adapted from *Ensembl genome browser*). B, Representative sequencing of the MDGA2 locus after successful insertion of the Myc tag in the Myc-MDGA2 KI colony founder mouse. C, Predicted translation of the MDGA2 locus after successful insertion of the Myc tag.

**Fig S3.** A, Representative low magnification confocal (left) and high magnification fluorescence tomography images taken in the SR (right) showing immunofluorescence labeling for DAPI, HA-MDGA1, and either glutamatergic (Homer1b/c, top panels) or GABAergic (NLGN2, bottom panels) postsynaptic markers in P15 HA-MDGA1 KI hippocampi. B, Quantification of the colocalization of dense labeled HA-MDGA1-ir and Homer1b/c-ir or NLGN2-ir postsynaptic marker puncta in the SR of WT and HA-MDGA1 KI P15 mice. The proportion of colocalized HA puncta is higher in KIs than WTs ($p \leq 0.001$ in both cases). For KI brains, the proportion of HA-MDGA1 puncta colocalized with Homer1b/c is higher than with NLGN2 ($p < 0.001$). C, D, In the HA-MDGA1 /Homer1b/c colocalization experiment, the number of HA-ir puncta was higher in KI samples ($p = 0.0188$, C), but the number of Homer1b/c-ir puncta was not different ($p = 0.2324$, D). E, F, In the HA-MDGA1/NLGN2 colocalization experiment, the number of HA-ir puncta was higher in KI samples ($p = 0.0315$, E), but the number of NLGN2-ir puncta was not different ($p = 0.6875$, F). G, Low magnification confocal (left) and high magnification fluorescence tomography image taken in the SR (right) showing immunofluorescence labeling for DAPI, HA-MDGA1, and glutamatergic (VGlut1, top panels) or GABAergic (VGat, bottom panels) presynaptic markers in P15 HA-MDGA1 KI hippocampi. H, Quantification of the colocalization of dense labeled HA-MDGA1-ir and VGlut1-ir or VGat-ir puncta in the SR of WT and KI P15 mice. The proportion of colocalized HA puncta is higher in KIs than WTs ($p = 0.0058$ for VGlut1, $p = 0.0250$ for VGat). For KI brains, the proportion of HA-MDGA1 puncta colocalized with VGlut1 is higher than with VGat (p<0.0020). I, J, In the HA-MDGA1/VGlut1 colocalization experiment, numbers of HA-ir and VGlut-ir puncta were not significantly different between genotypes (p=0.0610, I; p=0.9098, J). K, L, in the HA-MDGA1/VGat colocalization experiment, the number of HA-ir puncta was higher in KI samples (p=0.0346, K), but the number of VGat-ir puncta was not different (p=0.5039, L). Bar graphs represent mean ± SEM. n=3-5 mice per group of either sex. n.s., non-statistically significant; *, p<0.05; **, p<0.01; ***, p<0.001, Two-way ANOVA followed Tukey's post hoc test in B and H and unpaired T-test in C-F and I-L. Scale bars: Left panels: 200 µm. Right panels: 2 µm.

**Fig S4. MDGA2 can interact with GluN2A and GluN2B independently of GluN1.** Co-IP of overexpressed Flag-tagged GluN2A or GluN2B subunit of the NMDAR together or not with the obligatory GluN1 subunit the with HA-tagged MDGA2 in heterologous HEK cells. Quantification is shown to the right. Both GluN2A and GluN2B interact with MDGA2 independently of GluN1 presence (p=0.230 and p=0.370, respectively). n=3. n.s., non-statistically significant (Student's T-test). "WB", Western blot; "Ab", antibody.

**Fig S5: Knockdown of MDGA family effects spine structure**. A, Experimental timeline. B, Sample images using super-resolution structured illumination microscopy (SIM) of primary apical dendrites from neurons expressing GFP (n=10 neurons, 180 spines) or shRNAs against MDGA1,2 (n=10 neurons, 164 spines). Scale bar 5 µm. C, Spine density on primary apical dendrites (p > 0.05). D, Spine head diameter (p > 0.05). E, Spine neck length (p < 0.0001). F, Spine neck diameter (p < 0.0001). n.s., non-statistically significant; ***, p<0.001 (Student's T-test).

**Fig S6: Expression of MDGA1 decreases excitatory currents**. A, Immunoblot analysis of pCAG-HA-MDGAs (high expression) or pCAG-IRES-HA-MDGA (low expression) plasmids. WB, western blot. Ab, antibody. B, Total IRES-HA-MDGA1 lysate levels (means ± s.e.m.) compared to pCAG-HA-MDGA1 levels (p=0.0100, n=3) and total IRES-HA-MDGA2 lysate levels (means ± s.e.m.) compared to pCAG-HA-MDGA2 levels (p=0.0068, n=3). C, AMPAR (p=0.0015, n=12)- and NMDAR (p=0.0034, n=12)-mediated EPSC scatter plots displaying reductions in pCAG-HA-MDGA1 expressing cells compared to control cells. Open circles are individual pairs, filled circle is mean ± s.e.m. Black sample traces are control, green are transfected neurons. Scale bar denotes 25 pA and 0.1 s. Bar graphs plots transfected amplitude normalized to control cell ± s.e.m. D, AMPAR (p=0.0025, n=12)- and NMDAR (p=0.0038, n = 11)-mediated EPSC scatter plots displaying reductions in pCAG-IRES-HA-MDGA1 expressing cells compared to control cells. Bar graph, sample traces, and scale bar as in C. E, AMPAR (p=0.0020, n=11)- and NMDAR (p>0.0039, n=9)-mediated EPSC scatter plots displaying reductions in pCAG-HA-MDGA1 FY147-148AA expressing cells compared to control cells. Bar graph, sample traces, and scale bar as in c. OE, overexpressing. *, p<0.05; **, p<0.01

**Fig S7: Expression of MDGA1 decreases inhibitory currents in a neuroligin-dependent manner**. A, Scatter plots showing a reduction in IPSC-mediated currents in MDGA1 expressing cells compared to untransfected control cells (p=0.0023, n=14). Open circles are individual pairs, filled circle is mean ± s.e.m. Black sample traces are control, green are transfected neurons. Scale bar denotes 100 pA and 0.05 s. Bar graphs plots transfected amplitude normalized to control cell ± s.e.m. B, Scatter plots showing a reduction in IPSC-mediated currents in IRES-MDGA1 expressing cells compared to untransfected control cells (p=0.0078, n=8). Bar graph, sample traces, and scale bar as in A. C, Scatter plots showing no significant difference in IPSC-mediated currents in MDGA1 FY147-148 expressing cells compared to untransfected control cells (p>0.05, n=14). Bar graph, sample traces, and scale bar as in a. IPSC amplitudes recorded at 0 mV. OE, overexpressing.
